# A SMAD1/5-YAP signaling module drives radial glia self-amplification and growth of the developing cerebral cortex

**DOI:** 10.1101/558486

**Authors:** Sonia Najas, Isabel Pijuan, Anna Esteve-Codina, Susana Usieto, Juan D. Martinez, An Zwijsen, Maria L. Arbonés, Elisa Martí, Gwenvael Le Dréau

## Abstract

The growth and evolutionary expansion of the cerebral cortex are defined by the spatial-temporal production of neurons, which itself depends on the decision of radial glial cells (RGCs) to self-amplify or to switch to neurogenic divisions. The mechanisms regulating these RGC fate decisions are still incompletely understood. Here we describe a novel and evolutionarily conserved role of the canonical BMP transcription factors SMAD1/5 in controlling neurogenesis and growth during corticogenesis. Reducing the expression of both SMAD1 and SMAD5 in neural progenitors at early mouse cortical development caused microcephaly and an increased production of early-born cortical neurons at the expense of late-born ones, which correlated with the premature differentiation and depletion of the pool of cortical progenitors. Gain- and loss-of-function experiments performed during early cortical neurogenesis in the chick revealed that SMAD1/5 activity supports self-amplifying RGC divisions and restrain the neurogenic ones. Furthermore, we demonstrate that SMAD1/5 stimulate RGC self-amplification through the positive post-transcriptional regulation of the Hippo signaling effector YAP. We anticipate this SMAD1/5-YAP signaling module to be fundamental in controlling growth and evolution of the amniote cerebral cortex.

## Introduction

The cerebral cortex is the region of the human brain responsible for higher cognitive functions. Errors during its formation provoke a vast array of brain disorders that affect intellectual ability and social behaviour (Hu et al., 2014; Jayaraman et al., 2018). This highlights the relevance of identifying the mechanisms that govern cortical development, in particular those controlling its growth and the production of neurons from cortical stem and progenitor cells.

Emerging in the dorsal part of the telencephalon (pallium), the mammalian cerebral cortex first consists of a pseudo-stratified epithelial layer (also called the ventricular zone, VZ) formed by primitive neural stem cells, the neuroepithelial cells, that rapidly mature into radial glial cells (RGCs) (Taverna et al., 2014). Like neuroepithelial cells, RGCs contact both the ventricle and the basal lamina, they divide at the apical membrane and they can expand through self-amplifying divisions that produce two daughter RGCs (De Juan Romero and Borrell, 2015; Taverna et al., 2014). Neurogenesis is initiated when RGCs start producing daughter cells committed to the neuronal lineage. Neurogenesis can occur directly, when a RGC divides asymmetrically to generate another RGC and a daughter cell that differentiates directly into a neuron (De Juan Romero and Borrell, 2015; Taverna et al., 2014). Alternatively, neurogenesis can be indirect, whereby a RGC gives rise to a RGC and a basal progenitor (BP) that harbours a restricted lineage potential, delaminates from the VZ and divides basally. These BPs will increase the neuronal output, possibly self-amplifying for several rounds of divisions before generating two neurons through a final self-consuming division (Lui et al., 2011). While intermediate progenitor cells (IPCs) possess a very limited self-amplifying potential and represent the vast majority of cortical BPs in lissencephalic species, basal (or outer) RGCs harbour a considerable self-amplifying potential and they are responsible for the tremendous production of neurons and the larger neocortex observed in gyrencephalic species (Lui et al., 2011; Shitamukai et al., 2011; Wang et al., 2011). Therefore, the decision of a RGC to self-amplify or to give rise to neurons, either directly or indirectly, has a huge impact on the final number of neurons in the cerebral cortex.

As highlighted by the gene mutations causing primary microcephaly, a variety of intracellular events regulate cerebral cortical size, including centrosome behaviour and centriole biogenesis, DNA replication and repair, cytokinesis and apical-basal polarity (Jayaraman et al., 2018; Saade et al., 2018). Moreover, the balance between RGC self-amplification and neurogenesis appears to be regulated by extrinsic cues emanating from the ventricular fluid, meninges, blood vessels and neighbouring cells (Llinares-Benadero and Borrell, 2019; Martynoga et al., 2012; Taverna et al., 2014). Thus, the molecular events regulating RGC fate are complex and they are still not fully understood.

Seminal studies performed on cortical progenitors *in vitro* suggested that Bone morphogenetic protein (BMP) signaling regulates their neurogenic potential (Li et al., 1998; Mabie et al., 1999). The microcephaly described in *Bmp7* mutant mice, and the over-proliferation and premature differentiation reported in the brains of transgenic mice expressing constitutively active forms of the type-1 BMP receptors *Alk3/Bmpr1a* or *Alk6/Bmpr1b* supported this idea (Panchision et al., 2001; Segklia et al., 2012). However, definitive proofs of an instructive role for BMP signaling in cortical RGC fate decision *in vivo* are still lacking.

Here we found that the activity of the transcription factors SMAD1/5, two canonical effectors of BMP signalling, promotes cortical RGC self-amplification in both chick and mouse embryos, preventing their premature neurogenic switch and exhaustion, and thereby ensuring the appropriate production of the distinct classes of cortical excitatory neurons. In searching for the effectors of SMAD1/5 activity, we show that this role depends on the post-transcriptional regulation of YAP, a core component of the Hippo signaling pathway known to regulate cell growth and organ size. Together, our findings reveal an instructive and evolutionarily conserved role of the canonical BMP effectors SMAD1/5 in the control of RGC self-amplification in the developing cerebral cortex.

## Results

### SMAD1/5 activity is required for brain growth and cortical neurogenesis in mouse

To determine the role played by the canonical BMP pathway during mammalian cerebral cortex development, we focused on the function of its canonical effectors SMAD1/5/8 during mouse corticogenesis. *In situ* hybridization revealed that *mSmad1* and *mSmad5* transcripts are expressed throughout the rostral-caudal axis of the developing mouse cerebral cortex at embryonic day (E) 14.5 and are particularly enriched in the VZ, whereas *mSmad8* transcripts were not detected (Fig. 1A). In agreement with previous observations (Saxena et al., 2018), immunostaining at mid-corticogenesis with an antibody that specifically recognizes the active, carboxy-terminal phosphorylated form of SMAD1/5/8 (pSMAD1/5/8) revealed activity of these canonical BMP effectors in differentiating neurons as well as in both apically- and basally-dividing cortical progenitors (Fig. 1B). When quantified in pH3^+^ mitotic nuclei, pSMAD1/5/8 immunostaining revealed SMAD1/5 activity to be stronger in apically-dividing mouse RGCs than in basally-dividing IPCs (Fig. 1C), suggesting a correlation between high SMAD1/5 activity and RGC maintenance.

**Figure 1:**
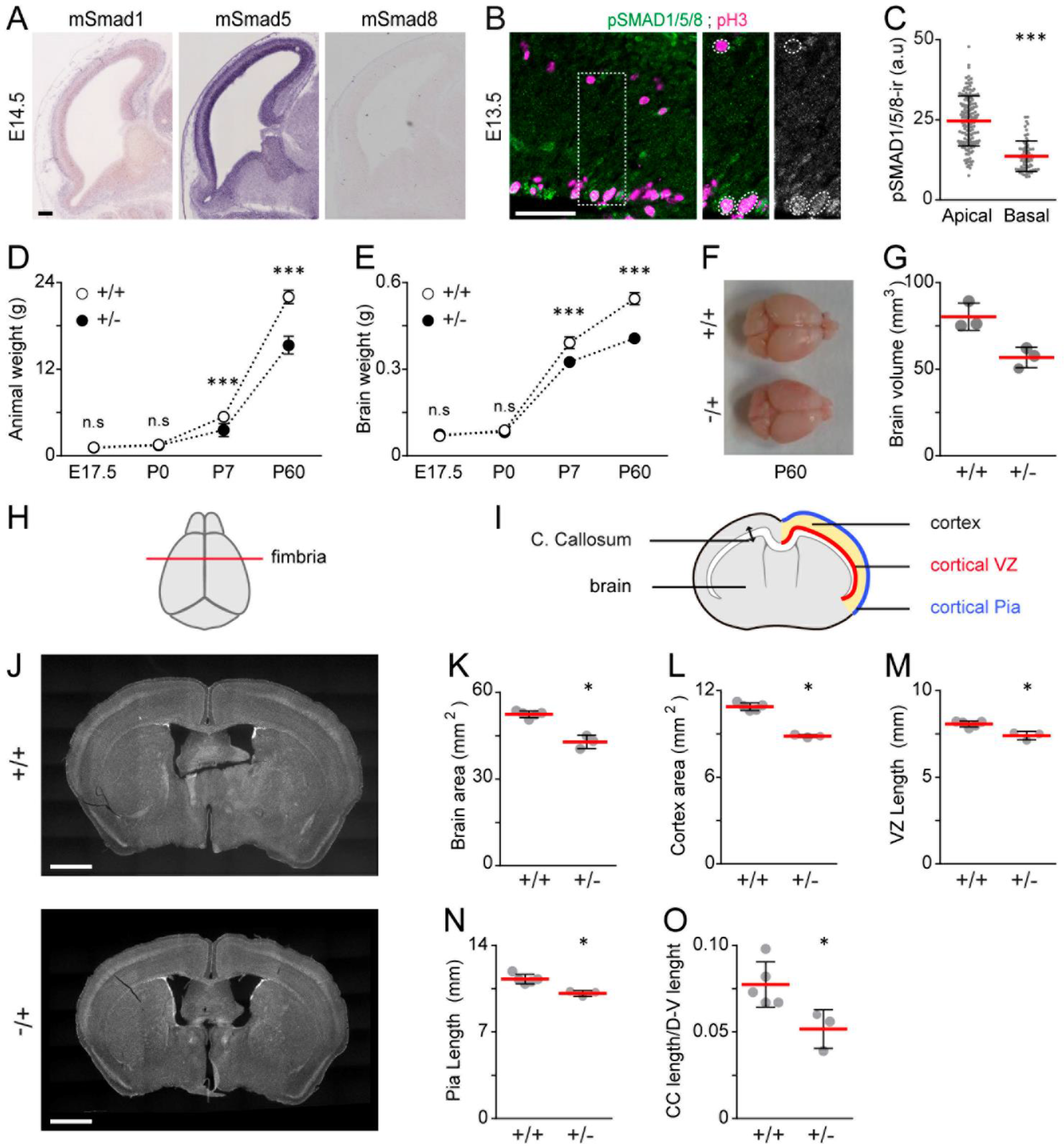
The SmadNes mutant mice present severe microcephaly and growth retardation. (A) Sagittal sections of the developing mouse dorsal telencephalon showing *mSmad1, mSmad5* and *mSmad8* transcripts at E14.5, obtained from Genepaint (https://gp3.mpg.de). **(B)** The pSMAD1/5/8 immunoreactivity at E13.5 and (C) its mean intensity ± s.d measured in 144 apical and 64 basal mitoses obtained from 3 mouse E13.5 embryos. (D) Weight ± s.d of the SmadNes mutant mice (*Smad1wt/fl;Smad5wt/fl;Nestin:Cre+/0*, +/−) and their control littermates *(Smad1^wt/fl^;Smad5^wt/fl^;Nestin:Cre^0/0^*, +/+) calculated from 17, 10, 12, 5 +/+ and 27, 9, 5, 3 +/− animals at E17.5, P0, P7 and P60, respectively. (E) Brain weight ± s.d calculated from 6, 10, 5, 5 +/+ and 16, 9, 3, 3 +/− animals at E17.5, P0, P7 and P60, respectively. (F) Dorsal view and (G) mean volume ± s.d of the brain of the SmadNes mutants and their control littermates at P60, obtained from 3 +/+ and 3 +/− animals. (H) Rostral-caudal brain level at which (I) the distinct parameters were analysed at P60. (J) Coronal sections obtained at the fimbria level (as shown in H) and the corresponding mean area ± s.d of (K) the whole brain and of (L) the cerebral cortex, the length ± s.d of (M) the cortical VZ and of (N) the cortical pia and (O) the thickness of the corpus callosum relative to whole dorsal-ventral brain thickness, all obtained from 5 +/+ and 3 +/− animals. Significance was assessed with the non-parametric Mann–Whitney test (C, K-O) or a two-way ANOVA + Sidak’s test (D, E). *P<0.05, ***P<0.001, ns: P>0.05. Scale bars, 50 μm (A, B), 250 μm (J). See also Figures S1 and S2.

To understand the role played by SMAD1/5 during mouse cerebral cortex development, we crossed *Smad1^fl/fl^;Smad5^fl/fl^* mice (Moya et al., 2012) with a *Nestin:cre* transgenic line that produces Cre-mediated recombination in neural progenitor cells and somites as early as E8.5 (Petersen et al., 2002), earlier than the more commonly used *Nestin:cre* transgenic line that produces efficient Cre-mediated recombination in cortical progenitors from mid-embryogenesis (Liang et al., 2012). In agreement with the early Cre-mediated recombination expected from this *Nestin:cre* transgenic line (Petersen et al., 2002), SMAD1 and SMAD5 protein levels were reduced by 54% and 32% in telencephalic extracts from E11.5 *Smad1^wt/fl^;Smad5^wt/fl^;Nestin:Cre^+/0^* heterozygous embryos relative to their control *Smad1^wt/fl^;Smad5^wt/fl^;Nestin:Cre^0/0^* littermates (Fig. S1). These *Smad1^wt/fl^;Smad5^wt/fl^;Nestin:Cre^+/0^* heterozygous mutant mice were viable and born following Mendelian ratios but were sterile, precluding the study of the homozygous compound mutants. Compared to their *Smad1^wt/fl^;Smad5^wt/fl^;Nestin:Cre^0/0^* littermates (hereafter referred to as controls or +/+), the *Smad1^wt/fl^;Smad5^wt/fl^;Nestin:Cre^+/0^* heterozygous mutant mice (hereafter referred to as SmadNes mutants or +/−) presented an overall growth retardation, including a reduction in brain weight, detected from postnatal day P7 (Fig. 1D, E). The brain of the adult (P60) SmadNes mutants showed a 25% reduction in weight (+/−: 0.407g ± 0.006 *vs* +/+: 0.544g ± 0.021; Fig.1E), that correlated to a 29% decrease in volume (Fig.1F,G) and a 18% reduction of its surface area in coronal sections (+/−: 42.91 mm^2^ ± 2.302 *vs* +/+: 52.46 mm^2^ ± 1.18, Fig. 1H-K). These reductions in brain weight and size were both greater than 3SD implying, according to clinical standards (Passemard et al., 2013), that the SmadNes heterozygous mutants suffer a severe microcephaly. The coronal areas of the whole brain and cerebral cortex were similarly reduced in adult SmadNes mutants (18% and 19% respectively; Fig.1K, L), and this reduction was constant across the rostral-caudal axis (Fig. S2). Constant decreases in the length of the cortical VZ and pia along this axis were also observed in the SmadNes mutant brain (Fig.1M, N and Fig.S2). Apart from these growth defects and a thinner corpus callosum (Fig. 1O), the brain of the SmadNes mutants did not present any major neuro-anatomical defects (Fig.1J and S2B), thereby suggesting that SMAD1/5 inhibition impaired growth equally in all brain regions.

The cerebral cortices of the adult SmadNes mutants and their control littermates showed comparable thicknesses and presented similar densities of NeuN^+^ neurons (Fig.2A-C). They also presented similar densities of macroglial cells, including SOX9^+^ astrocytes (Fig. S3A,B; Sun et al., 2017), and oligodendroglial cells (SOX10^+^, or OLIG2^+^;CC1^+^ oligodendrocytes and OLIG2^+^;CC1^-^ progenitors, Fig.S3C-F; Bhat et al., 1996; Claus Stolt et al., 2002). Taken altogether, these findings suggested that SMAD1/5 inhibition affects the tangential growth of the brain and the generation of its radial columns rather than its radial growth and the number of cells per radial unit. Nevertheless, we observed that the relative proportions of the different neuronal layers forming the cerebral cortex were altered in the adult SmadNes mutant (Fig.2A,C). The number of early-born neurons forming the deep layer L6 and its thickness were increased, whereas these parameters were diminished in the superficial L2/3 layer containing late-born callosal projection neurons (Fig.2A,C,D), consistent with the reduced thickness of the corpus callosum (Fig.1O). This phenotype was observed at the early postnatal stage P7 (Fig.2E-G), supporting the idea that these cortical defects originated during the embryonic phase.

**Figure 2:**
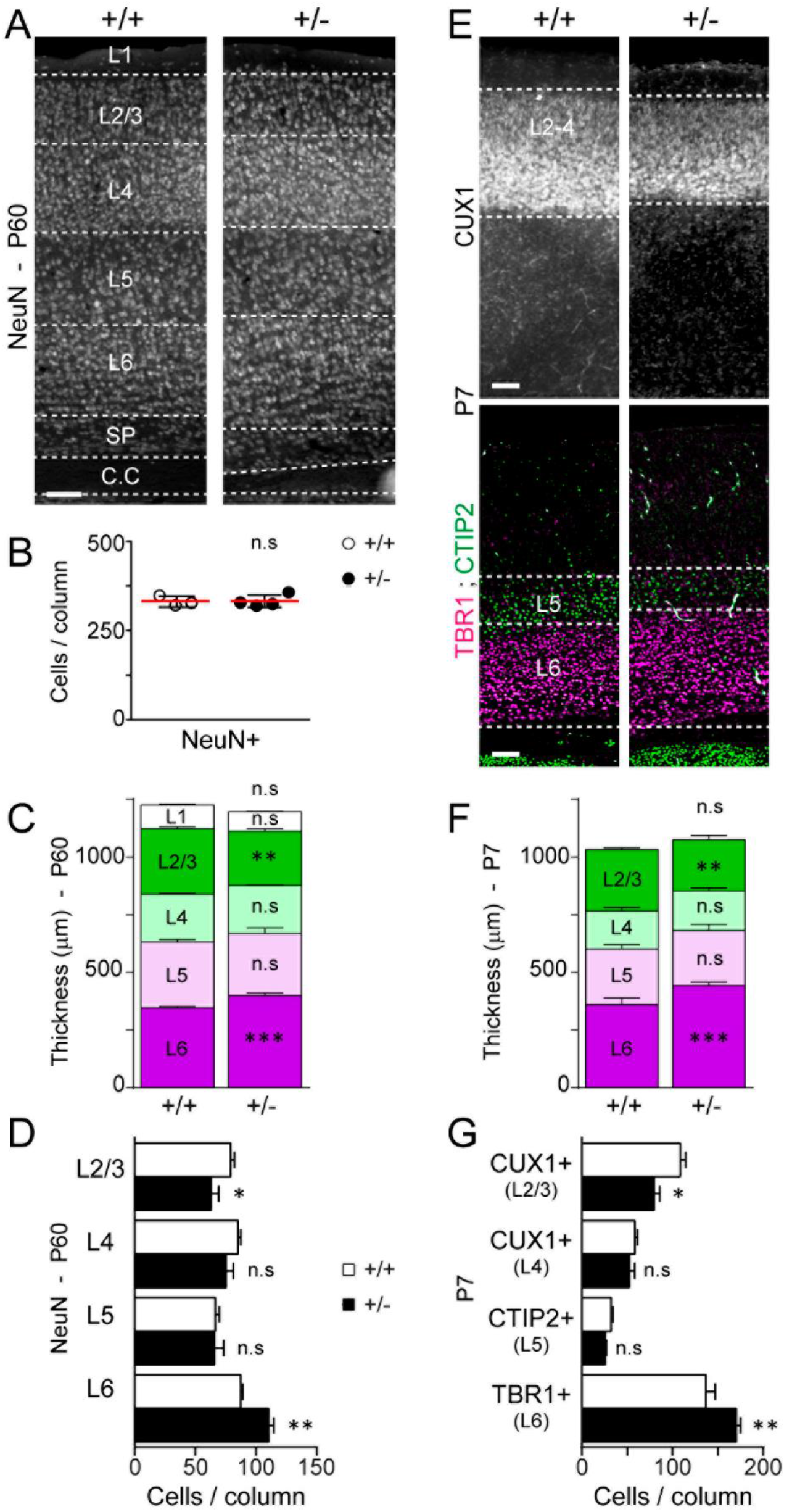
Inhibiting SMAD1/5 activity in mouse neural progenitors causes an increase in early-born neocortical neurons at the expense of late-born ones. (A) NeuN^+^ neurons present in coronal sections of the brains of SmadNes mutant mice (*Smad1^wt/fl^;Smad5^wt/fl^;Nestin:Cre^+/0^*, +/−) and their control littermates (*Smad1^wt/fl^;Smad5^wt/fl^;Nestin:Cre^0/0^*, +/+) at P60, and (B) their mean number ± s.d quantified in a 100 μm-wide cortical area, obtained from 3 +/+ and 4 +/− animals. (C) Mean thickness of the cortical neuronal layers ± s.e.m and (D) mean number of NeuN^+^ neurons ± s.e.m in the different layers in a 100 μm-wide cortical area at P60, obtained from 4 +/+ and 3 +/− animals. (E-G) Early- and late-born neocortical neurons present in coronal sections of the brains of SmadNes mutant pups and their control littermates at P7. (E) Early-born L6 (TBR1^+^), L5 (CTIP2^+^) and late-born L4-2/3 (CUX1^+^) projection neurons, (F) mean thickness of the layers ± s.e.m and (G) mean number of neurons ± s.e.m in the different layers in a 100 μm-wide cortical area, obtained from 5-7 +/+ and 5 +/− animals. Significance was assessed with the non-parametric Mann–Whitney test (B; C and F for the total cumulated thickness) or a two-way ANOVA + Sidak’s test (C, D, F, G). *P<0.05, **P<0.01, ***P<0.001, ns: P>0.05. Scale bars, 100 μm. See also Figure S3.

While the SmadNes mutant embryos did not present any obvious defect in telencephalic patterning (Fig.S4A), their programme of cortical neurogenesis did appear to be altered (Fig.3). Around the onset of cortical neurogenesis (E11.5), the SmadNes mutant embryos presented RGCs (PAX6^+^;TBR2^-^ cells) in correct numbers per radial area and of a normal cell size (Fig.3A-C and Fig.S4B). At this early neurogenic stage, the SmadNes mutant embryos also presented correct numbers of TBR2^+^ IPCs and a germinal zone of normal thickness (Fig.3B,D,E), yet their neuronal output was increased (Fig.3F,G). Accordingly, there were more early-born differentiating TBR1^+^ neurons in the SmadNes mutant cortex than in their control littermates from E11.5 onwards (Fig.3H,I; Bulfone et al., 1995). The developing cerebral cortex of the SmadNes mutant embryos contained fewer IPCs from E13.5 onwards and fewer RGCs at E17.5 (Fig.3B-D). This decrease in cortical progenitors at E17.5 was associated with a reduced thickness of the germinal zones (Fig.3E), a lower neuronal output (Fig.3F,G) and fewer late-born CUX1^+^ and SATB2^+^ neurons (Fig.3J,K and S5; Britanova et al., 2008; Nieto et al., 2004). Together, these data confirmed the developmental origin of the alterations in cortical projection neurons seen in the cerebral cortex of P7 and P60 SmadNes mutants (Fig.2). The mitotic indices of the cortical RGCs and IPCs were comparable in SmadNes mutants and controls at all stages examined (Fig.S6), suggesting that the proliferation rate of the cortical progenitors is not severely affected in SmadNes mutants. We thus reasoned that the increased production of early-born neurons, the premature reduction in the number of RGCs and IPCs and the decreased production of late-born neurons observed in the cerebral cortex of SmadNes heterozygous mutant embryos might likely be the result of a premature switch of RGCs from self-amplifying to neurogenic divisions during early corticogenesis.

**Figure 3:**
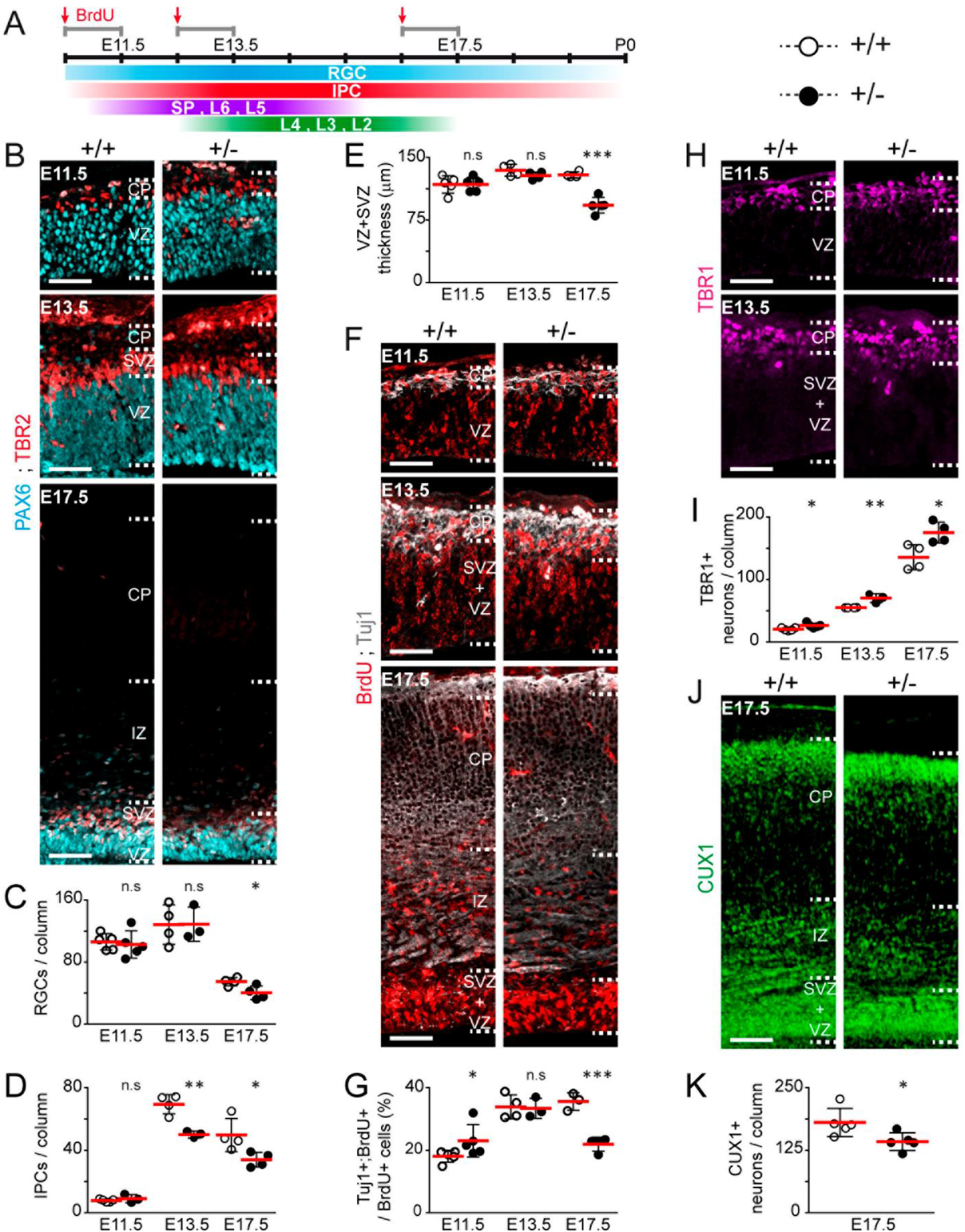
Inhibiting SMAD1/5 activity in mouse cortical progenitors causes premature neurogenesis and depletion of RGCs and IPCs. (A) Timeline of cortical neurogenesis and BrdU injections during mouse embryonic development. (B-J) Analysis of corticogenesis in the developing cerebral cortex of SmadNes mutant embryos (*Smad1^wt/fl^;Smad5^wt/fl^;Nestin:Cre^+/0^*, +/−) and their control littermates *(Smad1^wt/fl^;Smad5^wt/fl^;Nestin:Cre^0/0^*, +/+) at early (E11.5), mid (E13.5) and late (E17.5) embryonic stages. (B-D) Immunostaining of cortical progenitors and mean numbers ± s.d of (C) PAX6^+^;TBR2^-^ RGCs and (D) TBR2^+^ IPCs quantified in a 100 μm-wide cortical area, obtained from 5, 4, 4 +/+ and 5, 3, 4 +/− animals analysed at E11.5, E13.5 and E17.5, respectively. (E) Mean thickness of VZ+SVZ ± s.d, obtained from 5, 4, 4 +/+ and 5, 4, 5 +/− animals at E11.5, E13.5 and E17.5, respectively. (F, G) Neuronal output defined as the mean percentage ± s.d of BrdU^+^;Tuj1^+^/BrdU^+^ cells quantified in a 100 μm-wide cortical area 24 hours after a BrdU pulse (see A), obtained from 5, 4, 3 +/+ and 5,4,4 +/− animals at E11.5, E13.5 and E17.5, respectively. (H-K) Immunostaining and mean numbers ± s.d of early-born (TBR1, H, I) and late-born (CUX1; J, K) projections neurons quantified in a 100 μm-wide cortical area, obtained from (I) 5, 4, 4 +/+ and 5, 3, 4 +/− animals at E11.5, E13.5 and E17.5 and (K) 5 +/+ and 5 +/− animals at E17.5. Each dot represents the value of 1 animal. Significance was assessed with the two-sided unpaired t-test (C, D, G, I), the non-parametric Mann–Whitney test (K) or a two-way ANOVA + Sidak’s test (E). *P<0.05, **P<0.01, ***P<0.01, ns: P>0.05. Scale bars, 50 μm. CP: cortical plate, IZ: intermediate zone, SVZ: sub-ventricular zone, VZ: ventricular zone. See also Figures S4–S6.

### SMAD1/5 activity is required for RGC self-amplification during chick cortical neurogenesis

There is increasing evidence that the basic gene regulatory networks, progenitor cell types and cellular events governing the generation of neurons during corticogenesis are evolutionarily conserved between mammals and sauropsids, particularly birds (Cardenas et al., 2018; Le Dreau et al., 2018; Nomura et al., 2013; Suzuki et al., 2012; Yamashita et al., 2018). To assess whether SMAD1/5 regulate the mode of division of cortical RGCs, we turned to the developing chick cerebral cortex, as this avian model offered greater possibilities than the mouse to accurately manipulate SMAD1/5 activity at the onset of cortical neurogenesis.

The production of neurons in the developing chick cerebral cortex spans from E3 to E8 (Fig.4A,B; Suzuki et al., 2012). As in mammals, cortical neurogenesis in the chick is initiated by the onset of neurogenic divisions of PAX6^+^;TBR2^-^ RGCs that divide apically in the VZ (Fig.4A,B). From E5 onwards, the production of cortical neurons is enhanced by symmetric neurogenic divisions of basally-dividing TBR2^+^ IPCs (Fig.4A,B; Suzuki et al., 2012). *In situ* hybridization revealed that both *cSmad1* and *cSmad5* transcripts are expressed throughout the neurogenic period, mostly in the VZ where *cSmad8* transcripts were essentially absent (Fig.S7). Immunostaining with the pSMAD1/5/8 antibody at E5 revealed activity of SMAD1/5 in both apically- and basally-dividing cortical progenitors as well as in differentiating neurons (Fig.4C), as previously observed in the developing mouse cortex (Fig.1B). When quantified in pH3^+^ mitotic nuclei, SMAD1/5 activity was weaker in basal IPC divisions than in RGC divisions (Fig.4D). When quantified after *in ovo* electroporation of a pTis21:RFP reporter that is specifically activated during neurogenic divisions (Fig.S8; Le Dreau et al., 2014; Saade et al., 2013; Saade et al., 2017), SMAD1/5 activity was diminished in pTis21:RFP^+^ neurogenic divisions relative to pTis21:RFP^-^ self-amplifying divisions (Fig.4E,F). Therefore, a positive correlation exists between SMAD1/5 activity and the potential for RGC self-amplification during chick cortical neurogenesis.

**Figure 4:**
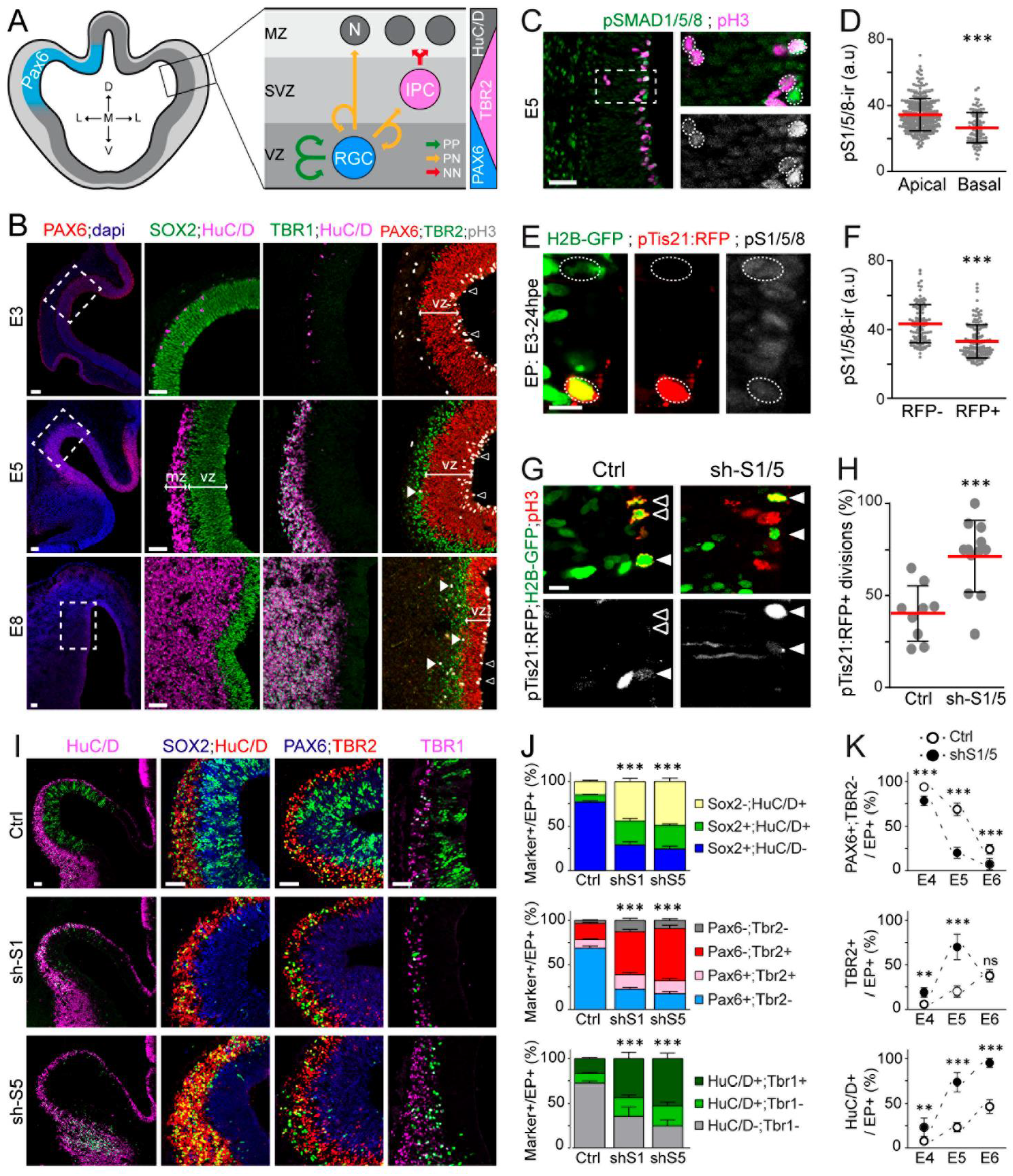
Inhibiting SMAD1/5 activity in chick cortical RGCs increases neurogenic divisions and causes their premature depletion and differentiation. (A) Cortical neurogenesis in the chick, its main cortical progenitor subtypes and their modes of division. (B) Coronal sections from 3, 5 and 8 days-old chick embryos showing the PAX6^+^ developing cerebral cortex, formed of SOX2^+^ neural progenitors, differentiating HuC/D^+^ and TBR1^+^ neurons, PAX6^+^;TBR2^-^ RGCs undergoing mitosis at the apical surface (black arrowheads) and TBR2^+^ IPCs dividing mostly basally (white arrowheads). (C) The active, phosphorylated form of SMAD1/5/8 (pSMAD1/5/8) immunoreactivity at E5, and (D) its mean intensity ± s.d measured in 276 apical and 93 basal mitoses obtained from 5 embryos. (E) The pSMAD1/5/8 immunoreactivity in mitotic cortical progenitors 24 hours after *in ovo* electroporation (IOE) with the pTis21:RFP reporter along with a control H2B-GFP-producing plasmid, and (F) its mean intensity ± s.d quantified in 137 pTis21:RFP^+^ and 107 pTis21:RFP^-^ divisions derived from 8 embryos. (G, H) The mean proportion ± s.d of electroporated (H2B-GFP+) cortical progenitors undergoing pTis21:RFP^+^ divisions (white arrowheads) after IOE of shRNAs targeting *cSmad1* or *cSmad5* (sh-S1/5, n=12 embryos) or their control (n=9). (I) Representative images and (J) mean proportions ± s.e.m of electroporated (H2B-GFP+) cells marked as (top) SOX2+/−;HuC/D+/−, (middle) PAX6+/−;TBR2+/− and (bottom) HuC/D+/−;TBR1+/−, assessed 48 hours after IOE with sh-S1 (n=8, 7, 5 embryos), sh-S5 (n=12, 5, 6) or their control (n=13, 8, 9). (K) The mean proportion ± s.d of electroporated cells identified as (top) PAX6+;TBR2-RGCs, (middle) TBR2+ committed cells and (bottom) HuC/D+ neurons, assessed 24 (E4), 48 (E5) and 72 (E6) hours after IOE and obtained from n ≥6 embryos per condition and stage. Significance was assessed with the non-parametric Mann–Whitney test (D, F), the two-sided unpaired t-test (H) or a two-way ANOVA + Tukey’s (J) or Sidak’s (K) test. **P<0.01, ***P<0.001, ns: P>0.05. Scale bars: 50 μm (B, C, I), 10 μm (E, G). VZ: ventricular zone, MZ: mantle zone. See also Figures S7 and S8.

Endogenous SMAD1/5 activity was inhibited from the onset of cortical neurogenesis by *in ovo* electroporation of sh-RNA plasmids that specifically target *cSmad1* or *cSmad5* and reduced their transcript levels by 40% and 60%, respectively (sh-S1/5; Le Dreau et al., 2012). Inhibiting SMAD1 or SMAD5 activity resulted in a similar phenotype, nearly doubling the proportion of electroporated RGCs undergoing neurogenic pTis21:RFP^+^ divisions (Fig.4G,H). This precocious switch to neurogenic divisions provoked a premature and accelerated depletion of electroporated PAX6^+^;TBR2^-^ RGCs, their accelerated progression towards a committed TBR2^+^ fate and ultimately, their differentiation into SOX2^-^;HuC/D^+^ and TBR1^+^ neurons (Fig.4I-K). Thus, full SMAD1/5 activity is required to support RGC self-amplification during chick cortical neurogenesis.

Conversely, enhancing SMAD5 activity through *in ovo* electroporation of a constitutively active SMAD5 mutant (SMAD5-S/D; Le Dreau et al., 2012) produced the opposite phenotype and reduced by 2 folds the proportion of electroporated RGCs undergoing neurogenic pTis21:RFP^+^ divisions (Fig.5A,B). This reduction in neurogenic divisions was associated with the electroporated cells remaining as PAX6^+^;TBR2^-^ RGCs and it impeded their transition into the neuronal lineage (Fig.5C,D). Notably, the SMAD5-S/D construct rescued the phenotype caused by sh-S5 (Fig.5E,F). In 5 out of 20 electroporated embryos, SMAD5-S/D electroporation itself caused the abnormal generation of ectopic rosettes of cortical progenitors, which developed an apical-basal polarity (Fig.S9). Together, these data revealed that the fine tuning of SMAD1/5 activity is required to properly balance the self-amplification of RGCs with the production of cortical excitatory neurons.

**Figure 5:**
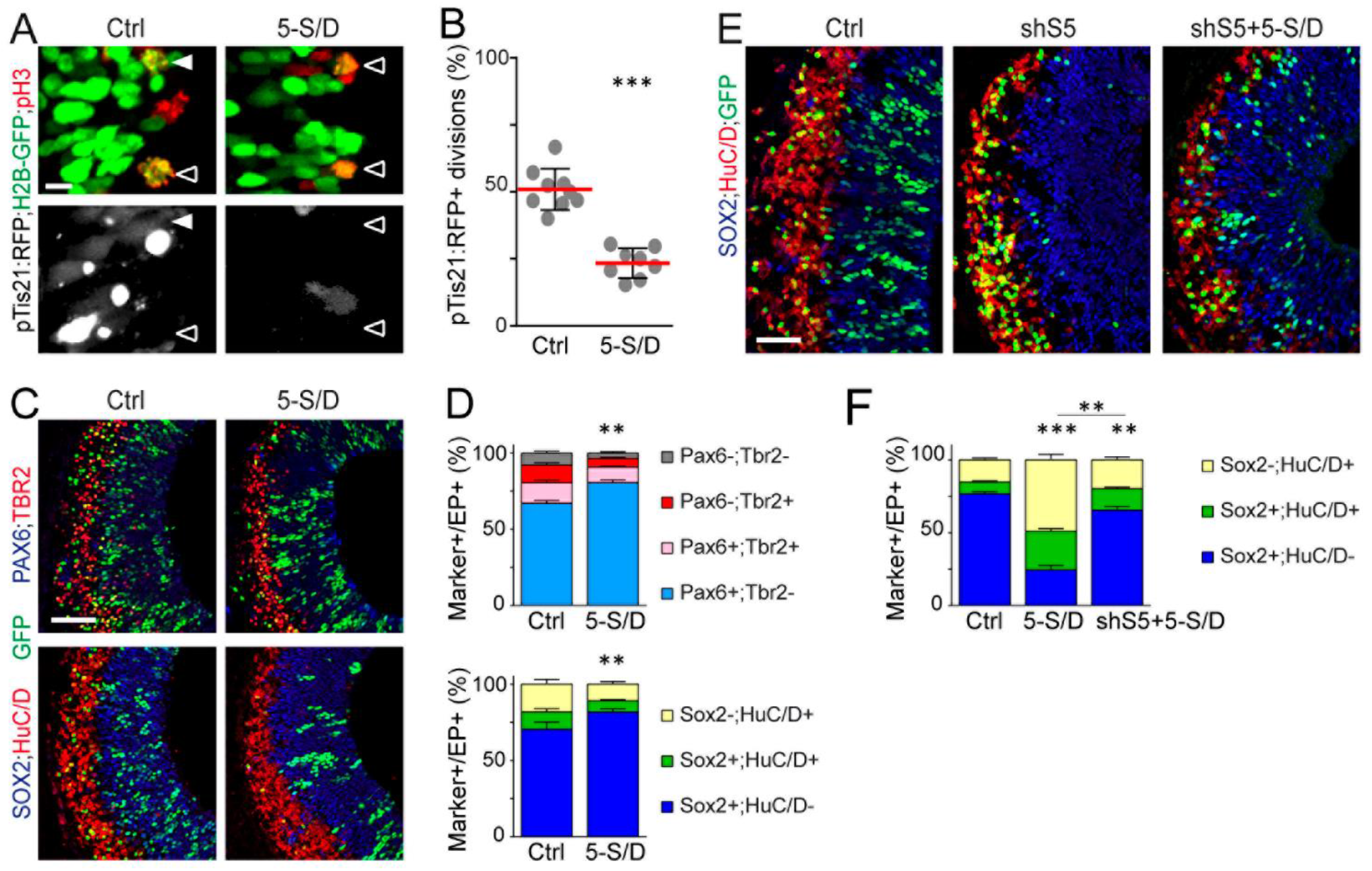
Increasing SMAD1/5 activity in chick cortical RGCs impedes neurogenic divisions and restrains differentiation. (A) Representative images and (B) mean proportion ± s.d of electroporated (H2B-GFP^+^) cortical progenitors undergoing pTis21:RFP^+^ divisions (white arrowheads) after IOE of a constitutively active SMAD5 mutant (5-S/D, n=8 embryos) or its control (n=9). (C) Representative images and (D) mean proportion ± s.e.m of electroporated (H2B-GFP^+^) cells marked as PAX6+/−;TBR2+/− and SOX2+/−;HuC/D+/− cells 48 hours after IOE of 5-S/D (n=15 and 14 embryos) or its control (n=7 and 10). (E) Representative images and (F) mean proportion ± s.e.m of of electroporated (H2B-GFP^+^) cells marked as SOX2+/−;HuC/D+/− cells 48 hours after IOE of a control plasmid (n=13), or sh-S5 combined with a control plasmid (sh-S5, n=12) or with the constitutively active SMAD5-S/D mutant (sh-S5+5-S/D, n=8). Significance was assessed with the two-sided unpaired t-test (B), a two-way ANOVA + Sidak’s (D) or + Tukey’s (F) tests. **P<0.01; ***P<0.001. Scale bars: 10 μm (A), 50 μm (C, E). See also Figures S9.

### SMAD1/5 regulate RGC self-amplification and early cortical neurogenesis through YAP

To identify the gene regulatory networks controlled by SMAD1/5 during cortical neurogenesis, cortical RGCs from SmadNes mutant and control E12.5 embryos were purified by FACS based on their Prominin1 expression (Corti et al., 2007), and their transcriptomes were compared by genome-wide RNA-sequencing (Fig.6A). A short list of differentially expressed transcripts (DETs) was identified (90 DETs with adj *p* < 0.05 and 128 with adj *p*<0.1: Fig.6B and Table S1). A gene ontology (GO) term enrichment analysis correlated this DET signature especially to biological processes related to the regulation of neurogenesis and cell biosynthesis (Fig.6C and Table S2), these two categories containing respectively 33 and 44 genes including 15 in common (Table S2). The genes retrieved in these two GO categories code for proteins playing various functions, the most frequent one being related to DNA-binding and the regulation of transcription (Fig.6D and Table S2). A Transfac/Jaspar analysis revealed that the promoter regions of these DETs are enriched in binding motifs for transcription factors of the TEAD and SP families (Fig.6E and Table S3). We considered particularly interesting these latter results pointing to an altered TEAD activity in response to SMAD1/5 inhibition, as the TEAD TFs indeed represent the transcriptional effectors of the Hippo signalling pathway, which regulates cell growth and organ size (Yu et al., 2015). Of the 42 DETs found to be related to either TEAD2 or TEAD4, 15 and 20 were retrieved in the GO terms related to neurogenesis and biosynthesis, respectively (Table S3). Using the genepaint database, we observed that the genes encoding these DETs are indeed expressed in the mouse developing cerebral cortex at mid-corticogenesis, mainly in the VZ (such as Camk2g, Cct3, Cd2bp2, Glce, Hmgn2, Phf21a, Spata13 and Trim24), the cortical plate (Ctnna2, Islr2, Nav1, Ndn and Reln) or in both (Agrn, Arid1a, Ehmt2, Hnrnpk, Klhl25, Spire1 and Slc25a51; Fig.S10).

**Figure 6:**
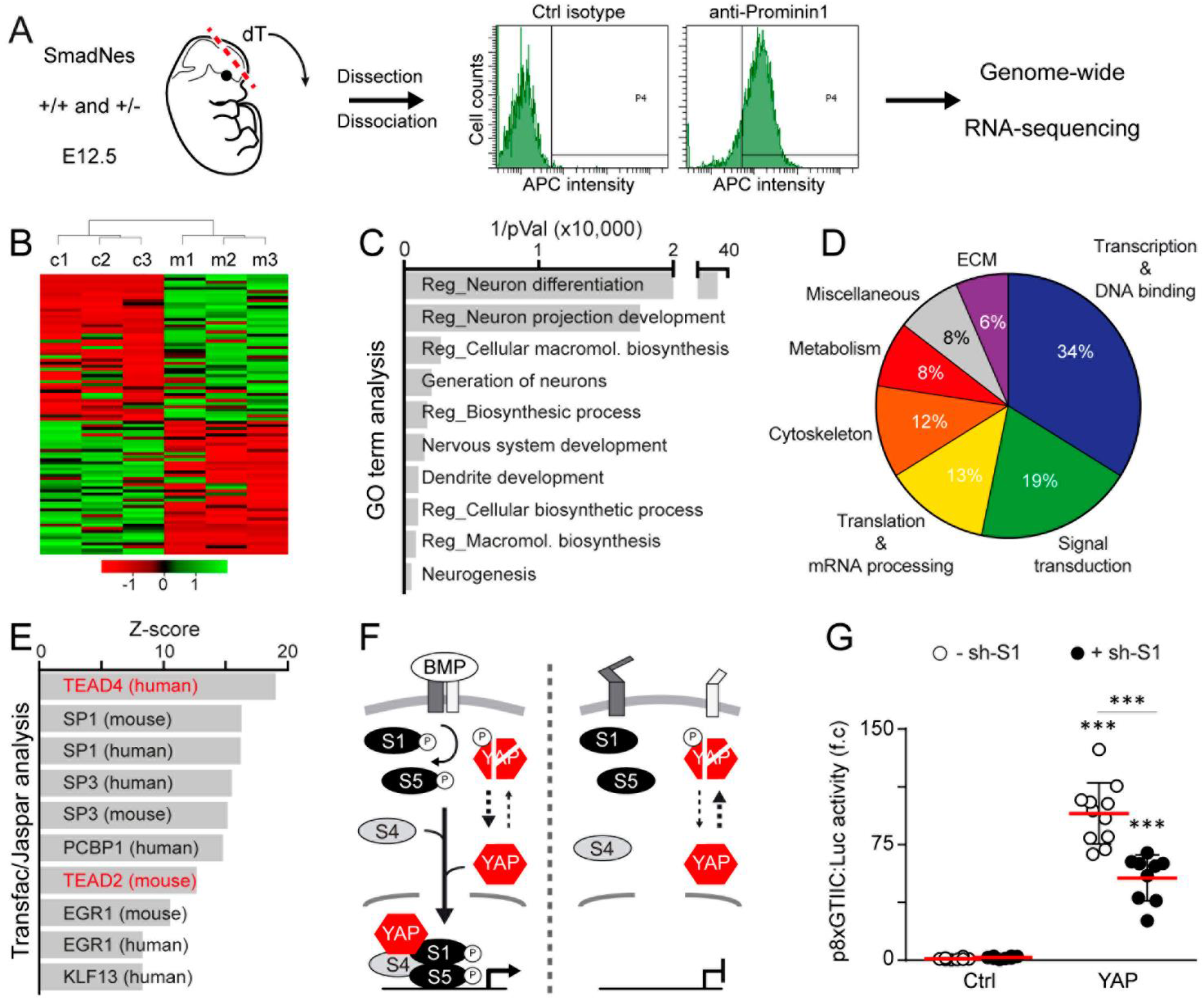
The transcriptional program regulated by SMAD1/5 in early mouse cortical RGCs correlates with an alteration of TEADs’ activity. (A) Methodology used to compare the transcriptome of cortical RGCs from E12.5 SmadNes mutant embryos (*Smad1^wt/fl^;Smad5^wt/fl^;Nestin:Cre^+/0^*, +/−) and control littermates *(Smad1^wt/fl^;Smad5^wt/fl^;Nestin:Cre^0/0^*, +/+) embryos. (B) Spearman rank correlation and heatmap representing the differentially expressed transcripts (DETs, with Adj *p*<0.05) between mutant and control cortical RGCs. (C) Top 10 gene ontology biological processes, (D) molecular function of the genes included in these biological processes and (E) Top 10 transcription factor binding motifs associated with the DETs (with adj *p*<0.1). (F) Upon BMP activity, SMAD1/5 are activated and can physically interact and act synergistically with YAP (left). When SMAD1/5 activity is reduced or abrogated, YAP is inactivated and primed for proteosomal degradation (right). (G) Mean activity ± s.d of a TEAD-responsive p8xGTIIC:luciferase reporter 24 hours after IOE of E3 chick telencephalons with a control plasmid (n=8 embryos), a wild type YAP1 construct (YAP, n=8), sh-S1 alone (n=11) or combined with YAP (n=9). Significance was assessed with a one-way ANOVA + Tukey’s test. ***P<0.001. See also Figure S10 and Tables S1-S4.

The activity of TEADs depends directly on the availability of their co-factors YAP/TAZ, which are themselves regulated by upstream kinases of the Hippo pathway (Yu et al., 2015). Cortical RGCs from the SmadNes embryos did not present any alteration in the transcript levels of the TEAD TFs, nor of other factors known to participate in Hippo signalling (Table S4). Thus, SMAD1/5 does not appear to regulate any member of the Hippo signalling pathway at the transcriptional level. However, previous findings established that YAP can physically interact with SMAD1/5 and that its activity is required for optimal SMAD1/5 activity in several cellular contexts, including cultured mouse embryonic stem cells, the drosophila wing imaginal disc and mouse astrocyte differentiation during postnatal development (Fig.6F; Alarcon et al., 2009; Huang et al., 2016). On the other hand, BMP-induced SMAD1/5 signalling has been reported to stimulate YAP protein stability, hence its activity (Fig.6F; Huang et al., 2016). We thus reasoned that YAP might participate together with SMAD1/5 in controlling RGC self-amplification during cortical neurogenesis. A luciferase assay performed after *in ovo* electroporation of a TEAD-responsive reporter (p8xGTIIC; Dupont et al., 2011) in the chick dorsal telencephalon confirmed that the ability of YAP overexpression to stimulate TEAD transcriptional activity was markedly impaired when SMAD1 activity was concomitantly inhibited (Fig.6G).

Therefore, we analysed YAP expression and activity relative to SMAD1/5 activity during cortical neurogenesis. In agreement with recent reports (Kostic et al., 2019; Saito et al., 2018), immunostaining for the YAP protein revealed that the active (nuclear) YAP was more intensely expressed in mitotic apical RGCs than in basally-dividing IPCs, and its expression was strongly correlated with SMAD1/5 activity during both chick and mouse cortical neurogenesis (Fig.7A-D and S11A-D). Inhibiting SMAD1/5 during chick cortical neurogenesis impaired YAP activity, as witnessed through the reduced nuclear YAP intensity in mitotic cortical RGCs and the global increase in the proportion of phosphorylated form of YAP (pYAP), which is primed for proteasomal degradation (Fig.7E,F and S11E,F). Similarly, the pYAP/YAP ratio increased in the cortex of E11.5 SmadNes mutant embryos, as shown by immunohistochemistry and western blotting (Fig.7G,H and S11G,H). These latter findings indicate that SMAD1/5 positively regulate YAP protein levels and activity both during mouse and chick cortical neurogenesis.

**Figure 7:**
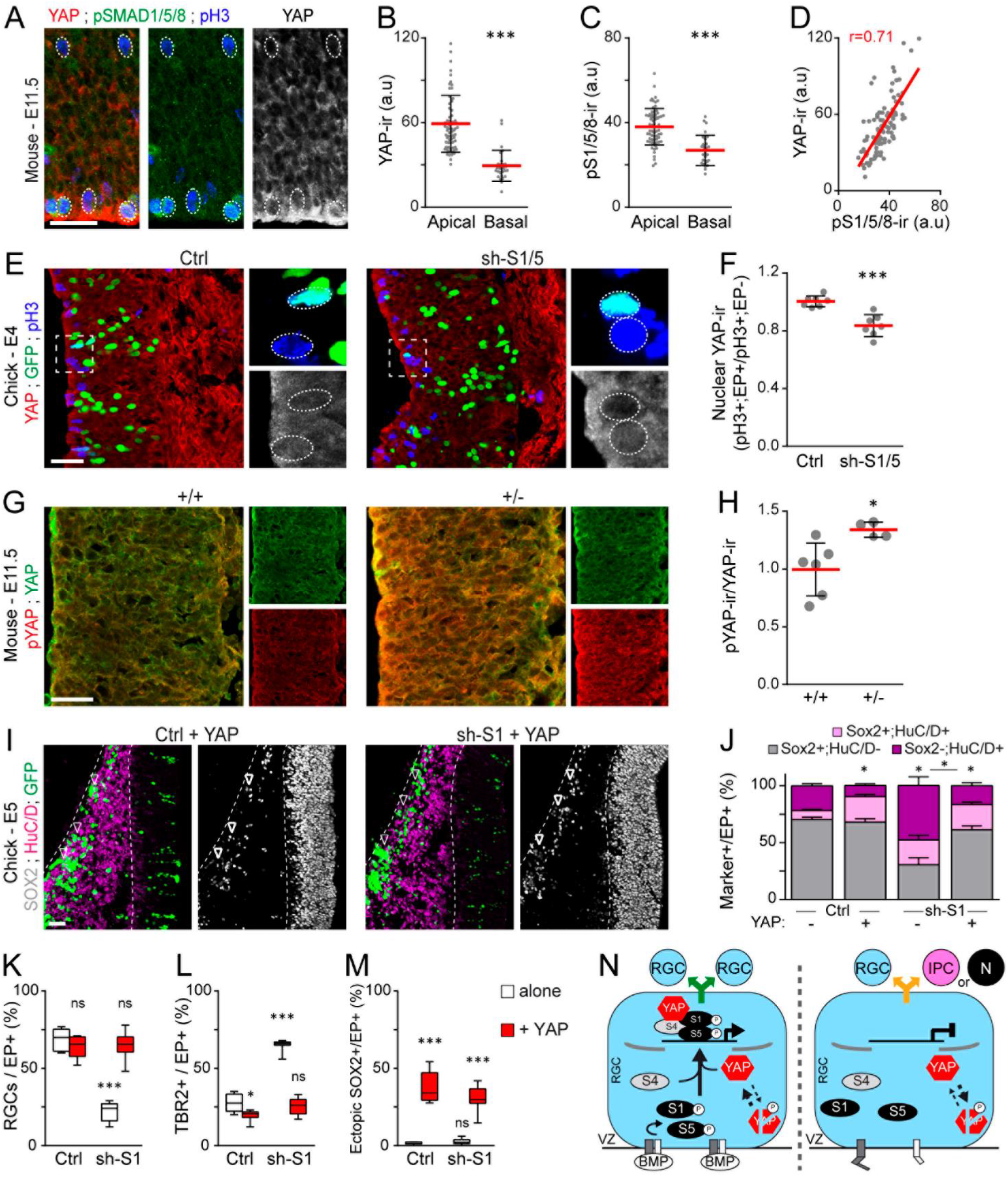
SMAD1/5 regulate early cortical neurogenesis through YAP. (A) YAP expression during early mouse corticogenesis (E11.5), relative to SMAD1/5 activity (pSMAD1/5/8). (B-D) The mean intensity ± s.d of (B) YAP and (C) pSMAD1/5/8 immunoreactivities, quantified in 70 apical and 28 basal pH3^+^ mitotic nuclei (n=3 embryos) and (D) their Pearson’s correlation coefficient r. (E) Total YAP immunostaining in the developing chick cerebral cortex and (F) its ratio ± s.d quantified in electroporated pH3^+^ mitotic cells relative to non-electroporated mitotic cells 24 hours after IOE with a control (Ctrl: 147 pH3^+^;GFP^+^ cells, 154 pH3^+^;GFP^-^ cells, n=7 embryos) or sh-S1/5 (sh-S1/5: 162 pH3^+^;GFP^+^ cells and 140 pH3^+^;GFP^-^ cells, n=7 embryos). (G) Immunostaining of pYAP and total YAP and (H) the quantification of the pYAP/YAP intensity ratio ± s.d in the developing cerebral cortex of E11.5 SmadNes mutants (*Smad1^wt/fl^;Smad5^wt/fl^;Nestin:Cre^+/0^*,+/−, n=4 embryos) and control littermates (*Smad1^wt/fl^;Smad5^wt/fl^;Nestin:Cre^0/0^,+/+,* n=4). (I) Representative sections and (J-M) mean proportion ± s.e.m of electroporated (GFP^+^) cells identified as (J) SOX2+/−;HuC/D+/−, (K) PAX6^+^;TBR2^-^ RGCs, (L) TBR2^+^ and (M) ectopic SOX2^+^ 48 hours after IOE of sh-S1 or its control electroporated alone (Ctrl, n=9 embryos; sh-S1, n=8) or together with a wild type YAP1 construct (Ctrl+YAP, n=8; sh-S1+YAP, n=8). (N) Model proposing that upon BMP signalling, the activated SMAD1/5 recruit YAP to promote RGC self-amplification. When SMAD1/5 activity is reduced or abrogated and YAP is inactivated and primed for degradation, their transcriptional activity is suppressed, enabling RGCs to undergo neurogenic divisions. Significance was assessed with the non-parametric Mann–Whitney test (B, C, F, H) or a two-way ANOVA + Tukey’s multiple comparisons test (J-M). *P<0.05, ***P<0.001, ns: P>0.05. Scale bars, 25 μm. See also Figure S11.

Finally, we tested whether increasing YAP activity could compensate for the phenotype caused by SMAD1/5 inhibition. *In ovo* electroporation of a wild type form of YAP rescued the premature exhaustion of RGCs driven by sh-S1, reverting it to control levels and impeding their progression towards differentiation into SOX2^-^;HuC/D^+^ neurons (Fig.7I-L). Intriguingly, YAP overexpression forced the vast majority of electroporated cells to remain SOX2^+^ (Fig.7I,J), with more than 30% being found ectopically in the mantle zone irrespective of SMAD1 inhibition (Fig.7I,M and S11I). Together, these results support a model whereby SMAD1/5 promotes RGC selfamplification and orchestrate cortical growth and neurogenesis through YAP (Fig.7N).

## Discussion

In this study, we identify a novel role for the canonical BMP effectors SMAD1/5 in the regulation of brain growth and cortical neurogenesis. More specifically, our findings demonstrated that SMAD1/5 activity stimulates cortical RGC self-amplification and impedes their premature switch to neurogenic divisions. By altering the balance between these modes of divisions, impairing SMAD1/5 activity during early corticogenesis has two main consequences.

First, the cerebral cortex of adult SmadNes mutant mice is smaller along the medial-lateral and rostral-caudal axes, although its thickness and the cell density are nearly normal. These observations suggest that reducing SMAD1/5 activity affects more severely the generation of radial columns than the number of cells per radial unit, emphasizing the importance of SMAD1/5 activity for cortical RGC self-amplification, which represents a critical step in the regulation of the growth of the cerebral cortex and which is particularly crucial before and during the early stages of cortical neurogenesis (Cardenas and Borrell, 2019). Of note, crossing the *Smad1^fl/fl^;Smad5^fl/fl^* mice with the *Nestin:cre* line from Jackson Laboratory, which produces an efficient Cre-mediated recombination in cortical progenitors only from late embryogenesis (around E17.5; Liang et al., 2012), did not cause any obvious alteration in cortical neurogenesis nor any apparent brain growth defects (Dr Eve Seuntjens, unpublished observations). These observations further support our conclusion that the role of SMAD1/5 in sustaining RGC self-amplification is especially crucial during the early steps of corticogenesis. Since the SmadNes heterozygous mutant mice present an overall reduction in brain size and weight, it is likely that SMAD1/5 play a similar role throughout the developing brain. We previously reported that SMAD1/5 promote neural progenitor self-amplification during spinal neurogenesis (Le Dreau et al., 2014; Le Dreau et al., 2018). Altogether, these data thus suggest that SMAD1/5 promote stem cell maintenance and growth throughout the developing CNS.

Second, the premature switch from RGC self-amplification to neurogenic divisions caused by SMAD1/5 inhibition altered the generation of the distinct classes of cortical projection neurons. Reducing SMAD1/5 activity during early mouse corticogenesis caused cortical progenitors to prematurely enter neurogenesis, enhancing the generation of early-born cortical neurons and prematurely exhausting the RGC and IPC pools,subsequently limiting the production of late-born cortical neurons. Related to this aspect, a recent study revealed that SMAD1/5 activity also orchestrates the transition from an early to late phase of neurogenesis during mouse cerebellum development, by repressing the late-born interneuron fate determinant Gsx1 (Ma et al., 2020). Importantly, our results strongly suggest that the dependence on SMAD1/5 activity to maintain RGC self-amplification and ensure appropriate neuronal production during corticogenesis is evolutionarily conserved in amniotes, at least between mammals and birds. This leads us to hypothesize that the transcription factors SMAD1/5 are part of the core ancestral gene regulatory network that governs corticogenesis throughout the amniote lineage, together with PAX6, the proneural bHLH proteins, the NOTCH and SLIT/ROBO signalling pathways (Cardenas et al., 2018; Le Dreau et al., 2018; Nomura et al., 2013; Suzuki et al., 2012; Yamashita et al., 2018).

The endogenous activity of SMAD1/5 was assessed using an antibody that specifically recognizes the active, carboxy-terminal phosphorylated form of SMAD1/5/8. This phosphorylation targeting the 3 carboxy-terminal serine residues is mediated by type-1 BMP receptors, whose activation depends on their interaction with type-2 BMP receptors, which itself depends on the binding of BMP ligands (Massague et al., 2005). Thus, the activity of SMAD1/5 described herein should reflect the activity of BMP ligands. Various members of the BMP family (including *Bmp2, Bmp4, Bmp5, Bmp6* and *Bmp7)* are expressed in the developing mouse cerebral cortex (Mehler et al., 1997). To our knowledge BMP7, whose deletion causes microcephaly in the mouse (Segklia et al., 2012), is the sole BMP ligand with a reported role in cortical neurogenesis. BMP7 is expressed by the hem, the meninges and the choroid plexus and can be detected in the cerebrospinal fluid (Segklia et al., 2012). BMP7 activity might thus be transduced to SMAD1/5 within RGCs through either their apical membrane or basal foot.

In agreement with previous studies (Alarcon et al., 2009; Saxena et al., 2018), the endogenous pSMAD1/5/8 immunoreactivity was particularly obvious in mitotic RGCs and IPCs. Although we do not rule out that the low levels of pSMAD1/5/8 immunoreactivity that we observed in other phases of the cell cycle might correspond to low levels of SMAD1/5 transcriptional activity, our data favour the idea that SMAD1/5 are activated just before or during mitosis. A growing body of literature reveals that many, if not most, of the genes transcribed during interphase are also transcribed during mitosis, albeit to low levels (Palozola et al., 2019). This low transcriptional activity occurring during mitosis, termed mitotic bookmarking, is believed to ensure transcriptional memory propagation from a mother cell to its daughters. Such activity has been observed for general promoter transcription factors, and for a growing list of tissue-specific transcription factors (Palozola et al., 2019). The detection of active pSMAD1/5 immunoreactivity in mitotic RGCs and IPCs observed herein thus supports the idea that SMAD1/5 could play a role in mitotic bookmarking.

Our RNA-seq approach correlated SMAD1/5 impairment with an altered activity of the Hippo signaling effectors TEADs. However, none of the TEAD family members nor other factors known to participate in Hippo signalling presented an altered transcriptional expression in mouse cortical RGCs in response to SMAD1/5 inhibition (Table S4), thereby pointing to a transcription-independent regulation of TEADs’ activity by SMAD1/5. To test if SMAD1/5 and TEADs might cooperatively regulate the same target genes, we analysed *in silico* the presence of SMAD1/5 binding motifs in the proximal promoter region (−2000bp≥TSS≥+500bp) of the genes retrieved from our RNA-seq that possess TEAD binding motifs. SMAD1/5 binding motifs were found only in 23% (6 out of 26) of the gene promoters containing TEAD2/4 binding motifs (Table S3). While these results suggest that SMAD1/5 and TEADs might not directly cooperate at the promoter level, it is worth mentioning that these two families of TFs appear to preferentially regulate transcription by binding to distal enhancers (Morikawa et al., 2011; Stein et al., 2015), thereby leaving the question about their direct cooperation open. By contrast, our findings clearly established that the functional cooperation between SMAD1/5 and YAP, a central actor in the Hippo signaling pathway (Yu et al., 2015), is crucial for the regulation of cortical growth and neurogenesis, whereby SMAD1/5 regulate YAP activity in early cortical RGCs. Accordingly, the tight regulation of YAP activity appears to be crucial for correct brain formation and growth. Cortical and general brain development is affected when YAP activity is enhanced, either directly or indirectly (Lavado et al., 2013; Lavado et al., 2018; Liu et al., 2018; Saito et al., 2018). More importantly, an aberrant increase in YAP activity has been linked to various types of cortical heterotopia, such as those observed in the Van Maldergem syndrome (Cappello et al., 2013; Liu et al., 2018). Our observation that YAP overexpression causes ribbon-like heterotopias of SOX2^+^ and PAX6^+^ progenitors in the developing chick cerebral cortex is therefore reminiscent of these severe mammalian cortical defects. Conversely, a reduction in YAP activity is plausibly one of the mechanistic events contributing to primordial dwarfism syndromes, whose defining features include microcephaly and general growth defects (Klingseisen and Jackson, 2011). Interestingly, the SmadNes heterozygous mutant mice present both microcephaly and growth retardation. Interestingly, the levels of YAP activity also determine the abundance and proliferative ability of neocortical basal progenitors (Kostic et al., 2019), such that its regulation might have contributed to the evolutionary diversification and expansion of the mammalian neocortex. Our findings suggest that the regulation of YAP activity by SMAD1/5 in RGCs might be evolutionarily conserved. It is thus tempting to speculate that modulations of the canonical BMP activity might have influenced the growth and expansion of the cerebral cortex during amniote evolution, a hypothesis that remains to be investigated further.

## Materials and Methods

### Animals

*Smad1^wt/fl^;Smad5^wt/fl^;Nestin:Cre^+/0^* embryos and postnatal mice were obtained by crossing *Smad1^fl/fl^;Smad5^fl/fl^* mice (Moya et al., 2012) with transgenic mice that express Cre-recombinase in neural progenitor cells and somites from E8.5 *(NesCre8* mice: Petersen et al., 2002). *Smad1^wt/fl^;Smad5^wt/fl^;Nestin:Cre^0/0^* littermates were used as controls. The days of the vaginal plug and birth were defined as E0.5 and P0, respectively. *Smad1^fl/fl^;Smad5^fl/fl^* and *NesCre8* mice were maintained in their original mixed genetic backgrounds (CD1, 129/ola, and C57BL6). All the experimental procedures were carried out in accordance with the European Union guidelines (Directive 2010/63/EU) and the followed protocols were approved by the ethics committee of the Parc Científic de Barcelona (PCB).

Fertilized white Leghorn chicken eggs were provided by Granja Gibert, rambla Regueral, S/N, 43850 Cambrils, Spain. Eggs were incubated in a humidified atmosphere at 38°C in a Javier Masalles 240N incubator for the appropriate duration and staged according to the method of Hamburger and Hamilton (HH: Hamburger and Hamilton, 1951). According to EU animal care guidelines, no IACUC approval was necessary to perform the experiments described herein, considering that the embryos used in this study were always harvested at early stages of embryonic development. Sex was not identified at these stages.

### In ovo electroporation

Unilateral *in ovo* electroporations were performed in the developing chick dorsal telencephalon at stage HH18 (E3, 69-72 hours of incubation). Analyses were performed specifically in the dorsal-medial-lateral region of the developing chick cerebral cortex to minimize any possible variability along the medial-lateral axis. Plasmids were diluted in RNAse-free water at the required concentration [0 to 4 μg/μl] and injected into the right cerebral ventricle using a fine glass needle. Electroporation was triggered by applying 5 pulses of 50 ms at 22.5 V with 50 ms intervals using an Intracel Dual Pulse (TSS10) electroporator. Electroporated chicken embryos were incubated back at 38°C and recovered at the times indicated.

### Plasmids

Inhibition of *cSmad1* and *cSmad5* expression was triggered by electroporation of short-hairpin constructs inserted into the pSuper (Oligoengine) or pSHIN vectors together with a control H2B-GFP-producing plasmid as previously reported (Kojima et al., 2004; Le Dreau et al., 2012; Le Dreau et al., 2018). Electroporation of 2-4 μg/μl of these constructs caused specific and reproducible 40% and 60% inhibition of the target expression (Le Dreau et al., 2012). The pCAGGS_SMAD5-SD_ires_GFP, its control pCAGGS_ires_GFP (pCIG), as well as the pTis21:RFP reporter used to assess the modes of divisions undergone by spinal progenitors, were previously described in details (Le Dreau et al., 2012; Megason and McMahon, 2002; Saade et al., 2013). The pCAGGS_Flag-YAP1_ires_GFP construct was obtained by subcloning from a pCDNA:Flag-YAP1 kindly provided by Conchi Estaras (Addgene plasmid #18881, deposited by Yosef Shaul; Levy et al., 2008). The TEAD-responsive p8xGTIIC:luciferase reporter was kindly provided by Sebastian Pons (Addgene plasmid #34615, deposited by Stefano Piccolo; Dupont et al., 2011).

### Luciferase assay

TEAD transcriptional activity was assessed following electroporation of the p8xGTIIC:luciferase reporter together with a renilla luciferase reporter used for normalization, in combination with the indicated plasmids required for experimental manipulation. Embryos were harvested 24 hours later, the electroporated telencephalic region carefully dissected and homogenized in a Passive Lysis Buffer on ice. Firefly- and renilla-luciferase activities were measured by the Dual Luciferase Reporter Assay System (Promega).

### In situ hybridization

Chicken embryos were recovered at the indicated stages, fixed overnight at 4 °C in 4% paraformaldehyde (PFA), rinsed in PBS and processed for whole mount RNA *in situ* hybridization following standard procedures. Probes against chick *Smad1* (#chEST899n18) and *Smad8* (#chEST222h17) were purchased from the chicken EST project (UK-HGMP RC). Probe against *cSmad5* was kindly provided by Dr Marian Ros. Hybridized embryos were post-fixed in 4% PFA and washed in PBT, and 45 μM-thick vibratome sections (VT1000S, Leica) were mounted and photographed under a microscope (DC300, Leica). The data show representative images obtained from 3 embryos for each stage and probe. The images of *mSmad1, mSmad5 and mSmad8* expression in the developing mouse dorsal telencephalon at E14.5 and those of candidate target genes selected from the RNAseq analysis were all obtained from the Genepaint database (https://gp3.mpg.de).

### Histology and Immunohistochemistry

Mouse embryos were recovered at the indicated stages and their heads fixed by immersion in 4% PFA for 24 hours at 4 °C, cryoprotected with 30% sucrose in PBS, embedded in Tissue-Tek O.C.T. (Sakura Finetek), frozen in isopentane at −30 °C and sectioned coronally on a cryostat (Leica). Cryosections (14 μm) were collected on Starfrost pre-coated slides (Knittel Glasser) and distributed serially. Postnatal and adult mice were deeply anesthetized in a CO_2_ chamber and trans-cardially perfused with 4% PFA. The brains were removed, post-fixed and vibratome (40 μm) sections were then distributed serially. Chicken embryos were carefully dissected, fixed for 2 hours at room temperature (RT) in 4% PFA, rinsed in PBS and cryoprotected with 30% sucrose in PBS and 16 μm-thick coronal sections prepared with a cryostat.

For both species, immunostaining was performed following standard procedures. After washing in PBS containing 0.1% Triton X-100 (PBT), the sections were blocked for 1 hour at RT in PBT supplemented with 10% bovine serum albumin (BSA). When necessary, sections were submitted to an antigen retrieval treatment before blocking by boiling sections for 10 min in sodium citrate buffer (2 mM citric acid monohydrate, 8 mM tri-sodium citrate dihydrate, pH 6.0). For BrdU immunostaining, sections were incubated before blocking in 50% formamide in 2X SSC at 64 °C for 10 min followed by an incubation in 2N HCl at 37 °C for 30 min and finally 10 min in 0.1 M boric acid (pH 8.5) at RT. Sections were then incubated overnight at 4 °C with the appropriate primary antibodies (Table S5) diluted in a solution of PBT supplemented with 10% BSA or sheep serum. After washing in PBT, sections were incubated for 2 hours at RT with the appropriate secondary antibodies diluted in PBT supplemented with 10% BSA or sheep serum. Alexa488-, Alexa568- and Cy5-conjugated secondary antibodies were obtained from Invitrogen and Jackson Laboratories. Sections were finally stained with 1 μg/ml DAPI and mounted in Mowiol (Sigma-Aldrich).

### Cell cycle exit assay

Pregnant female mice received an intra-peritoneal injection of BrdU (100 mg/kg; Sigma) and were sacrificed 24 hours later. Embryos were collected and processed as described above. Sections were immunostained for BrdU and Tuj1 and the cell cycle exit rate (neuronal output) was estimated by quantifying the proportion of BrdU^+^-immunolabelled cells that were Tuj1^+^ (% of Tuj1^+^;BrdU^+^/BrdU^+^ cells).

### Image acquisition and treatment

Optical sections of mouse and chick embryo fixed samples (coronal views) were acquired at RT with the Leica LAS software, in a Leica SP5 confocal microscope using 10x (dry HC PL APO, NA 0.40), 20x (dry HC PL APO, NA 0.70), 40x (oil HCX PL APO, NA 1.25-0.75) or 63x (oil HCX PL APO, NA 1.40-0.60) objective lenses. Maximal projections obtained from 2μm Z-stack images were processed in Photoshop CS5 (Adobe) or ImageJ for image merging, resizing and cell counting. Optical sections of postnatal and adult mouse samples were acquired with a Leica AF7000 motorized wide-field microscope. Cell counting in embryo, postnatal and adult mouse samples were performed in a 100 μm-wide column of the lateral cortical wall, as indicated in the figures. Cell counts were performed in a minimum of 3 sections of the same rostral-caudal level per embryo or postnatal mouse sample. Cell counting and measurements of the layers’ thickness in the P7 and P60 mouse cerebral cortex was done based on DAPI staining combined to, CUX1 (Nieto et al., 2004), CTIP2 (Arlotta et al., 2005), TBR1 (Bulfone et al., 1995) and NeuN immunostaining. Quantification of pSMAD1/5/8, PAX6, TBR2, YAP and pYAP intensities was assessed using the ImageJ software. Cell nuclei of mitotic pH3^+^ cells or H2B-GFP^+^ electroporated and neighboring non-electroporated cells were delimitated by polygonal selection, and the mean intensity was quantified as mean gray values.

### Brain morphometry

Morphometric parameters were estimated from serial coronal sections stained with DAPI, using the ImageJ software. Images were collected in a Leica AF7000 microscope using 5X or 10X objective. The brain volume was calculated according to the Cavalieri principle (Gundersen and Jensen, 1987), which consists in multiplying the distance between sections by their area, using a set of serial consecutive sections spanning from Bregma +1.10 mm to −1.82 mm (Paxinos and Franklin, 2001).

### Western blot

Total protein extracts (≈ 40 μg) were resolved by SDS-PAGE following standard procedures and transferred onto a nitrocellulose membrane (Hybond-ECL, Amersham Biosciences) that was probed with the primary antibodies (Table S5) whose binding was detected by infra-red fluorescence using the LI-COR Odyssey IR Imaging System V3.0 (LI-COR Biosciences).

### Purification of mouse Prominin1^+^ RGCs

The purification of Prominin1^+^ cortical RGCs was achieved by fluorescence-activated cell sorting (FACS) of dissociated cells obtained from the forebrain of *Smad1^wt/fl^;Smad5^wt/fl^;Nestin:Cre^+/0^* and *Smad1^wt/fl^;Smad5^wt/fl^;Nestin:Cre^0/0^* E12.5 embryos. Forebrains of littermates with the same genotype were pooled and incubated in Hanks’ balanced salt solution (HBSS) containing 0.6% Glucose and 5 mM EDTA for 5 min at 37 °C. Cells were mechanically dissociated by gentle trituration and collected by centrifugation at 300 g for 10 min at 4 °C. Cells were resuspended in 200 μl of incubation media (PBS containing 0.6% glucose, 2% foetal bovine serum and 0.02% NaN_3_) and kept for 10 min at 4 °C with mild agitation. Cells were then incubated for 30 min at 4 °C with an APC-conjugated anti-Prominin1 antibody (eBioscience, #17-1331-81) diluted at 0.2 mg/ml. Incubation with a rat IgG1-APC antibody was performed in parallel to define non-specific fluorescence. After the incubation, cells were centrifuged at 300 g for 10 min at 4 °C and incubated with 1 ml of incubation media. Dissociated cells were then filtered through a falcon tube with a cell strainer cap (BD Biosciences; BD falcon 12 x 75 mm) in which the filter was previously dampened with 500 μl of incubation media. The number of isolated cells was determined using a Neubauer chamber and cells were diluted to a final concentration of 2-3 x 10^6^ cells/ml. DAPI (20 ng/mL: Vector Labs) was added to identify dead cells. Sorting was performed with a BD FACS Aria™ Fusion (BD Biosciences) cytometer. After sorting, collector tubes were centrifuged at 1000 g for 10 min at 4 °C and cell pellets stored at −80 °C.

### RNA-sequencing

RNA was extracted from pools of 6 x 10^5^ FACS-isolated cells using the miRNeasy Micro Kit (Qiagen). Quantification of total RNA was performed by Qubit^®^ RNA BR Assay kit (Thermo Fisher Scientific) and RNA 6000 Nano Bioanalyzer 2100 Assay (Agilent) was used to estimate the total RNA integrity. The 3 pairs of samples of RNAs from SmadNes mutant and control RGCs presenting the best RNA integrity were used for sequencing. Sample preparation protocol for the RNASeq libraries was following the manufacturer’s recommendations of KAPA Stranded mRNA-Seq Illumina^®^ Platforms Kit (Roche-Kapa Biosystems). The libraries were sequenced on HiSeq2500 (Illumina) in paired-end mode with a read length of 2 x 100 bp using TruSeq SBS Kit v4 (Illumina). Each sample was sequenced in a fraction of a sequencing v4 flow cell lane, following the manufacturer’s protocol. Image analysis, base calling and quality scoring of the run were processed using the manufacturer’s software Real Time Analysis (RTA 1.18.66.3) and followed by generation of FASTQ sequence files. RNA-Seq paired-end reads were mapping against *Mus musculus* reference genome (GRCm38) using STAR version 2.5.3a with ENCODE parameters for long RNA. Isoforms were quantified using RSEM version 1.3.0 with default parameters for stranded sequencing and the gencode version M15. Differential isoform analysis was performed with DESeq2 version 1.18 with default parameters. We considered differentially expressed transcripts those showing a *p*-adjusted value < 0.05 or *p*-adjusted value <0.1 (extended list). Fold-change (FC) values between genotypes (SmadNes mutants over controls) are expressed in Log2 (Table S1). The GO term enrichment analysis (biologic process) of the extended DET signature was performed with the PANTHER classification system (http://pantherdb.org) and the Transfac/Jaspar analysis using Enrichr. The binding motifs for SMAD1 and SMAD5 were obtained from Jaspar. The proximal promoter region (−2000bp≥TSS≥+500bp) of each gene of interest was obtained from the USCS browser and the presence of the distinct TF binding motifs in the promoters was determined using the software FIMO (http://meme-suite.org/tools/fimo) (Table S3).

### Statistical analyses

No statistical method was used to predetermine sample size. The experiments were not randomized. The investigators were not blinded to allocation during experiments. Statistical analyses were performed using the GraphPad Prism 6 software (GraphPad Software, Inc.). Unless noted otherwise (see quantifications), cell counts were typically performed on 3-5 images per embryo and *n* values correspond to different embryos or animals. The normal distribution of the values was assessed by the Shapiro-Wilk normality test. Significance was then assessed with a two-sided unpaired t-test, one-way ANOVA + Tukey’s test or two-way ANOVA + Sidak’s or Tukey’s test for data-presenting a normal distribution, or alternatively with the non-parametric Mann– Whitney test for non-normally distributed data. The following convention was used: n.s: P>0.05; *P<0.05, **P<0.01, ***P<0.001. The detailed information related to quantifications are detailed in the figure legends.

## Acknowledgements

we thank the members of M.L.A’s and E.M’s laboratories for their discussion of this study. We thank E. Rebollo and the IBMB Molecular Imaging platform, J. Comas and the PCB Flow Cytometry facility, and the CNAG-CRG Sequencing Unit for their assistance. We are grateful to E.J. Robertson and W. Zhong for providing the *Smad1^fl/fl^* and *Nestin:cre* mice, to E. Seuntjens for sharing information, and to C. Estarás, S. Pons, M. Ros and M. Wegner for providing reagents.

## Author contributions

Conceptualization: S.N. and G.L.D.; Methodology: S.N., I.P., A.E-C., S.U., J.D.M. and G.L.D.; Investigation: S.N., I.P., A.E-C. and G.L.D.; Resources: A.Z., M.L.A and E.M; Visualization: G.L.D.; Writing - Original Draft: G.L.D.; Funding Acquisition: M.L.A. and E.M.; Supervision: M.L.A., E.M. and G.L.D.

## Data and materials availability

All data is available in the main text or supplemental information.

## Correspondence and requests for materials

should be addressed to G.L.D.

## Supplementary Information

### List of supplementary items

Figure S1: Reduction in SMAD1/5 protein levels in the SmadNes heterozygous mutant mouse.

Figure S2: The size reduction of the adult SmadNes mutant brain is constant across the rostral-caudal axis.

Figure S3: No alteration in the density of macroglial cells in the cerebral cortex of SmadNes mutant mice.

Figure S4: Absence of patterning and cell size defects in the cerebral cortex of SmadNes mutant embryos.

Figure S5: The number of SATB2+ neurons is decreased in the cerebral cortex of SmadNes mutant embryos.

Figure S6: The mitotic index of RGCs and IPCs is not altered in the cerebral cortex of SmadNes mutant embryos.

Figure S7: Expression of cSmad1/5/8 during chick cortical neurogenesis.

Figure S8: The activity of the pTis21:RFP reporter identifies neurogenic divisions during early chick cortical neurogenesis.

Figure S9: Increasing SMAD5 activity causes the abnormal generation of ectopic rosettes of cortical progenitors.

Figure S10: Gene expression pattern of RNAseq candidates in the mouse telencephalon.

Figure S11: SMAD1/5 regulate early cortical neurogenesis through YAP.

Table S1: List of differentially expressed transcripts between cortical RGCs from SmadNes mutant and control E12.5 embryos.

Table S2: Gene ontology (GO) term enrichment analysis.

Table S3: Transfac/Jaspar analysis and genes whose promoters are enriched in TEAD binding motifs.

Table S4: Expression of the genes of the Hippo signalling pathway in cortical RGCs from SmadNes mutant and control E12.5 embryos.

Table S5: List of primary antibodies used in this study.

---

**Figure S1:**
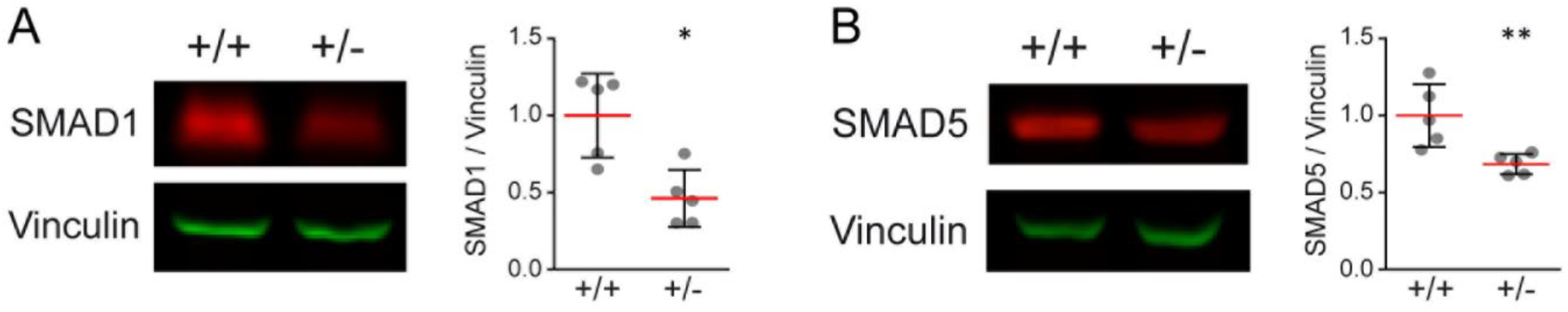
Reduction in SMAD1/5 protein levels in the SmadNes heterozygous mutant mouse. (A-B) The levels of SMAD1 (A) and SMAD5 (B) proteins were assessed by Western blot in extracts obtained from telencephalons of E11.5 SmadNes heterozygous mutant embryos *(Smad1^wt/fl^;Smad5^wt/fl^;Nestin:Cre^+/0^*, +/−) and their control littermates *(Smad1^wt/fl^;Smad5^wt/fl^;Nestin:Cre^0/0^,* +/+), relative to the levels of Vinculin. The data represent the mean ratio ± s.d obtained from 5 mutant embryos and 5 control littermates. Significance was assessed with the non-parametric Mann–Whitney test (A, B). *P<0.05, **P<0.01.

**Figure S2:**
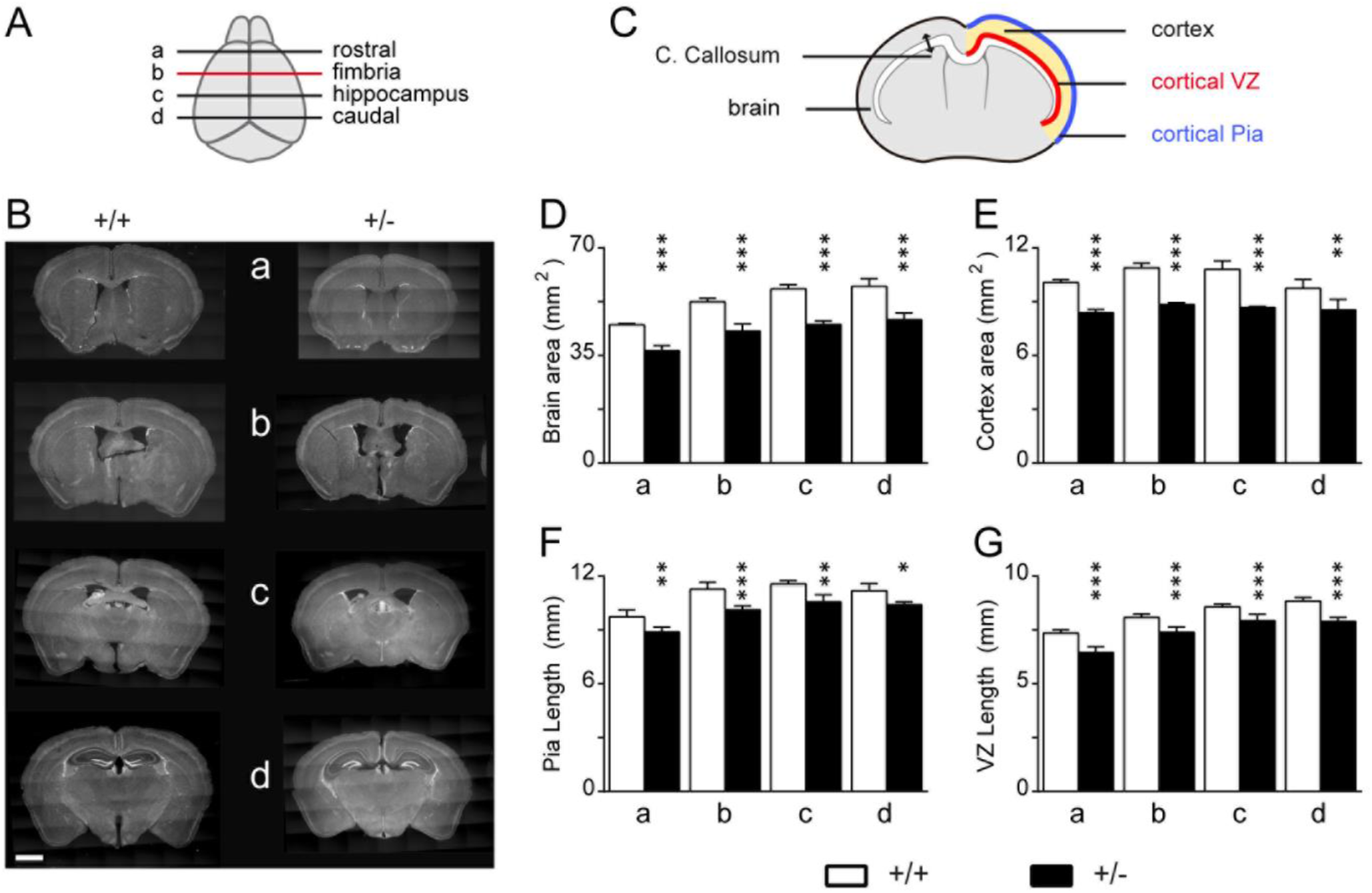
The size reduction of the adult SmadNes mutant brain is constant across the rostral-caudal axis. (A) Representative rostral-caudal levels and (B) coronal sections of the adult brain (P60) of the SmadNes mutant mice (*Smad1^wt/fl^;Smad5^wt/fl^;Nestin:Cre^+/0^*, +/−) and their control littermates *(Smad1^wt/fl^Smad5^wt/fl^;Nestin:Cre^0/0^*, +/+) in 4 different positions along the rostral-caudal axis (a to d, from rostral to caudal; b is also shown in Fig.1J), and (C) the distinct parameters analysed. (D,E) Mean area ± s.d of (D) the whole brain and of (E) the cerebral cortex. (F,G) Mean length ± s.d of (F) the cortical pia and of (G) the cortical VZ. n=5 embryos for +/+, n=3 for +/−. Significance was assessed with a two-way ANOVA + Sidak’s test (D-G). *P<0.05, ***P<0.01, ***P<0.001. Scale bar: 250 μm (B).

**Figure S3:**
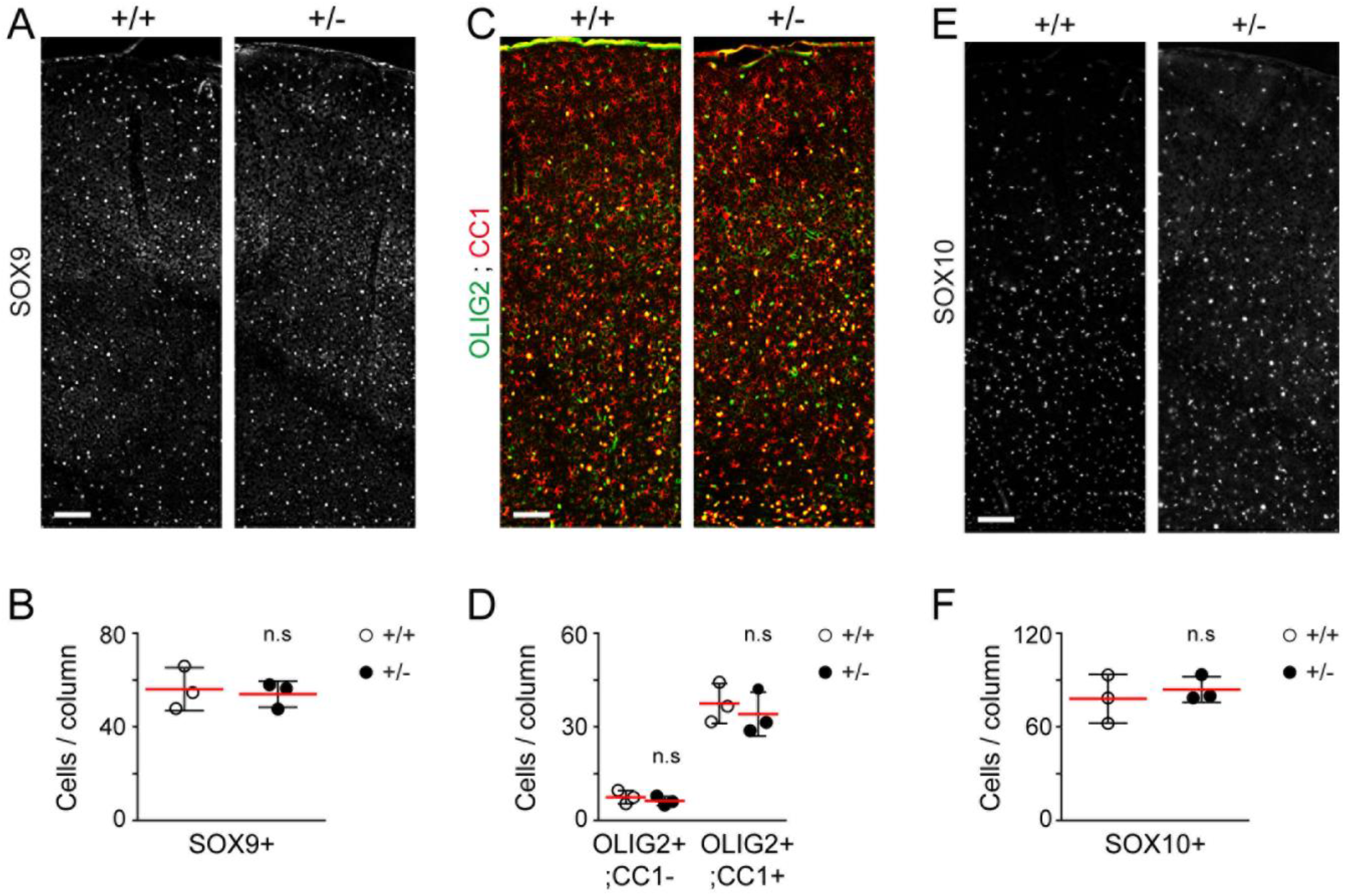
No alteration in the density of macroglial cells in the cerebral cortex of SmadNes mutant mice. The density of macroglial cells present in the brains of SmadNes mutant mice (*Smad1^wt/fl^;Smad5^wt/fl^;Nestin:Cre^+/0^*, +/−) and their control littermates (*Smad1^wt/fl^;Smad5^wt/fl^;Nestin:Cre^0/0^*, +/+) was assessed in coronal sections at P60. (A, B) SOX9^+^ astrocytes and (C-F) oligodendroglial cells identified as (C, D) OLIG2^+^;CC1^+^ oligodendrocytes and their OLIG2^+^;CC1^-^ progenitors or as (E, F) SOX10^+^ oligodendroglial cells, and (B, D, F) their mean number ± s.d quantified in a 100 μm-wide radial column, obtained from 3 +/+ and 3 +/− animals. Significance was assessed with the non-parametric Mann–Whitney test. ns: P>0.05. Scale bars, 100 μm.

**Figure S4:**
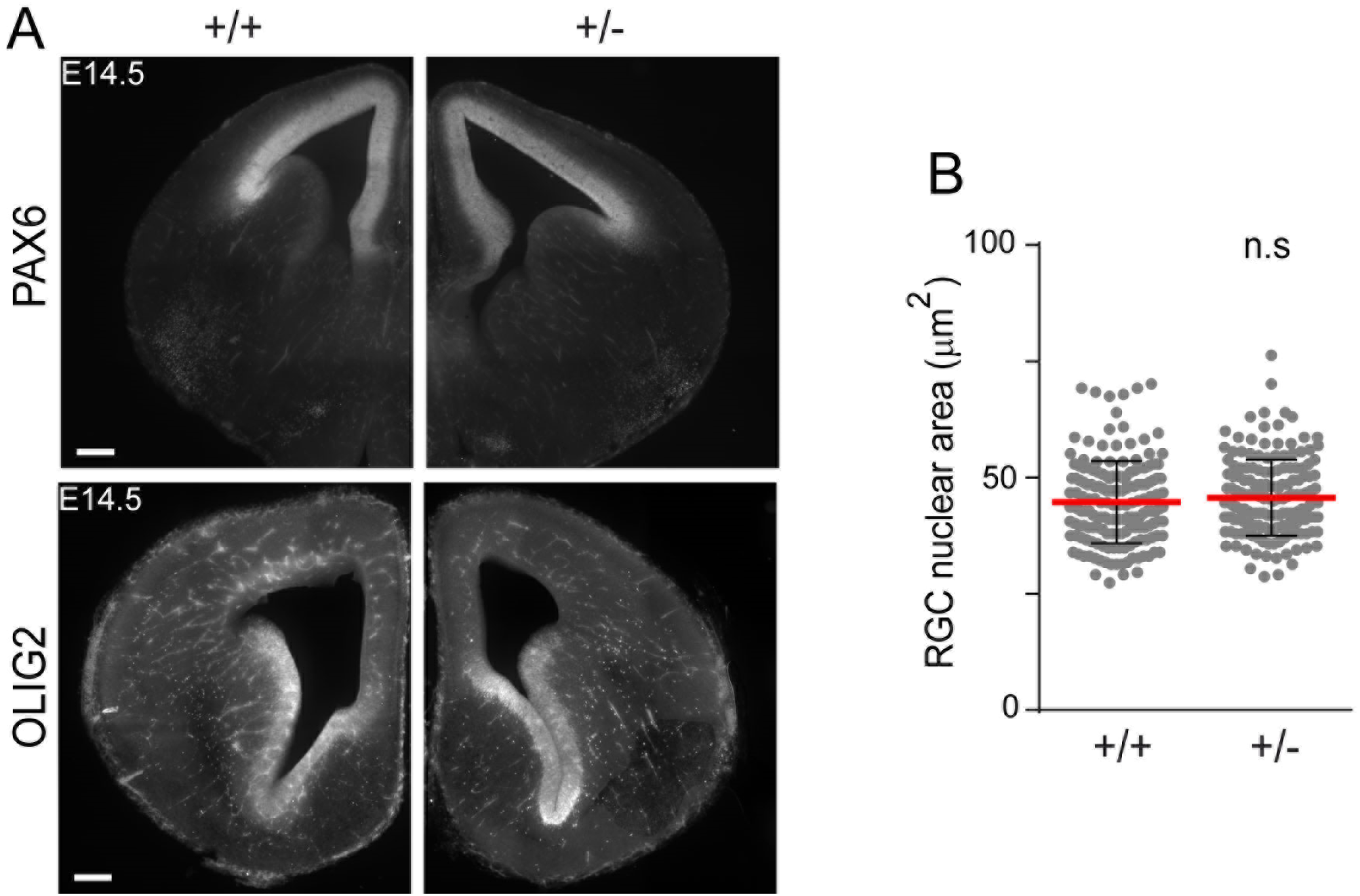
Absence of patterning and cell size defects in the cerebral cortex of SmadNes mutant embryos. (A) Coronal sections immunostained for the pallial and sub-pallial patterning markers PAX6 and OLIG2 in the telencephalon of E14.5 SmadNes mutant embryos (*Smad1^wt/fl^;Smad5^wt/fl^;Nestin:Cre^+/0^*, +/−) and their control littermates (*Smad1^wt/fl^;Smad5^wt/fl^;Nestin:Cre^0/0^*, +/+). (B) Mean nuclear area ± s.d of the PAX6+;TBR2-RGCs in the ventricular zone of E11.5 SmadNes mutant embryos and control littermates, quantified in 170 cells derived from 5 embryos for each genotype. Each dot represents the value of 1 cell. Significance was assessed with the non-parametric Mann–Whitney test (B). ns: P>0.05. Scale bars, 100 μm.

**Figure S5:**
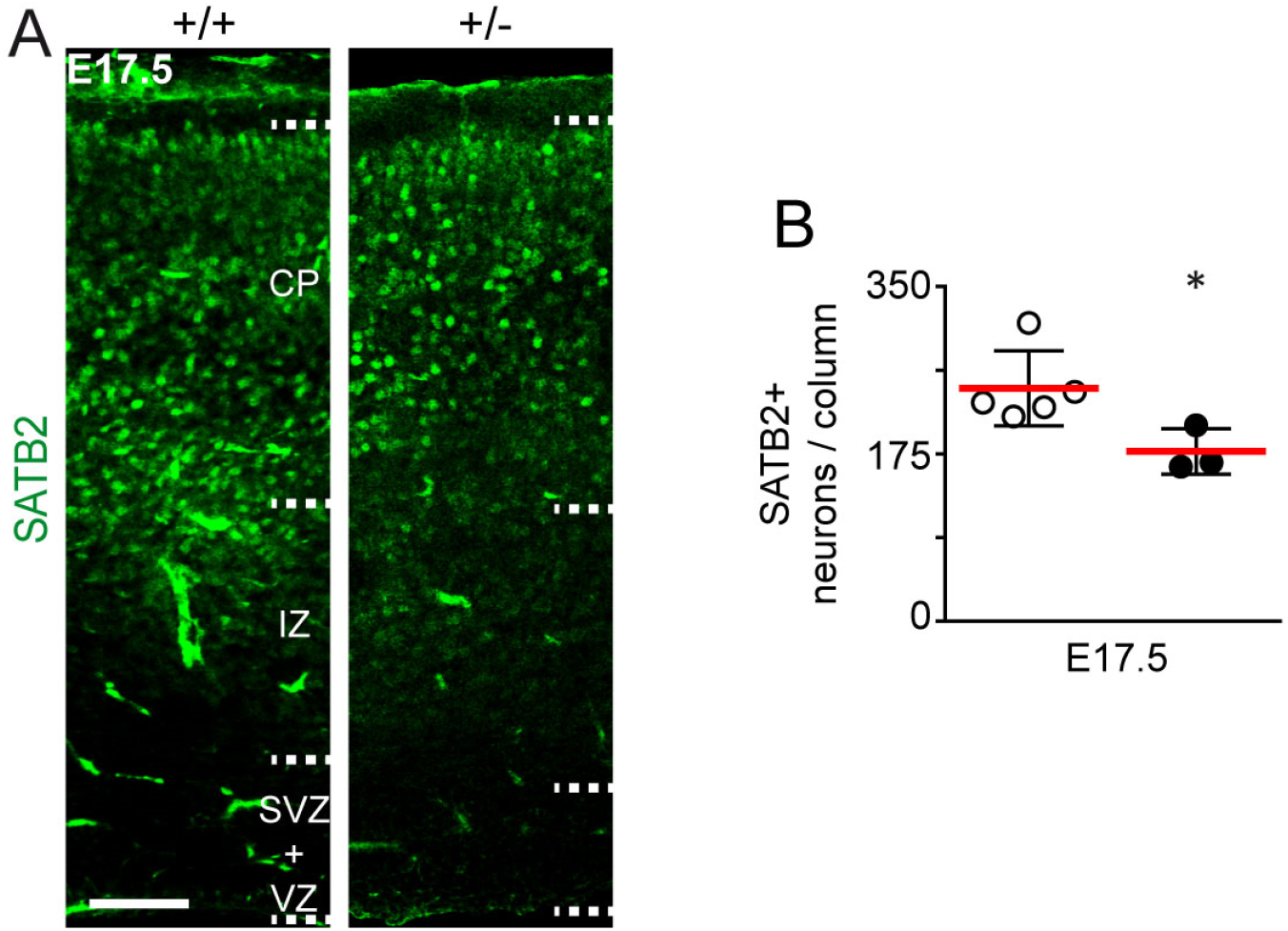
The number of SATB2+ neurons is decreased in the cerebral cortex of SmadNes mutant embryos. (A) Immunostaining for SATB2, which reveals mid- and late-born projections neurons, in the developing cerebral cortex of E17.5 SmadNes mutant embryos (*Smad1^wt/fl^;Smad5^wl/fl^;Nestin:Cre^+/0^’,* +/−) and control littermates *(Smad1^wt/fl^;Smad5^wt/fl^;Nestin:Cre^0/0^*, +/+). (B) Mean number of SATB2+ neurons ± s.d quantified in a 100 μm-wide cortical area, obtained from 5 +/+ and 3+/− embryos. Significance was assessed with the non-parametric Mann–Whitney test. *P<0.05. Scale bar, 50 μm. CP: cortical plate, IZ: intermediate zone, SVZ: sub-ventricular zone, VZ: ventricular zone.

**Figure S6:**
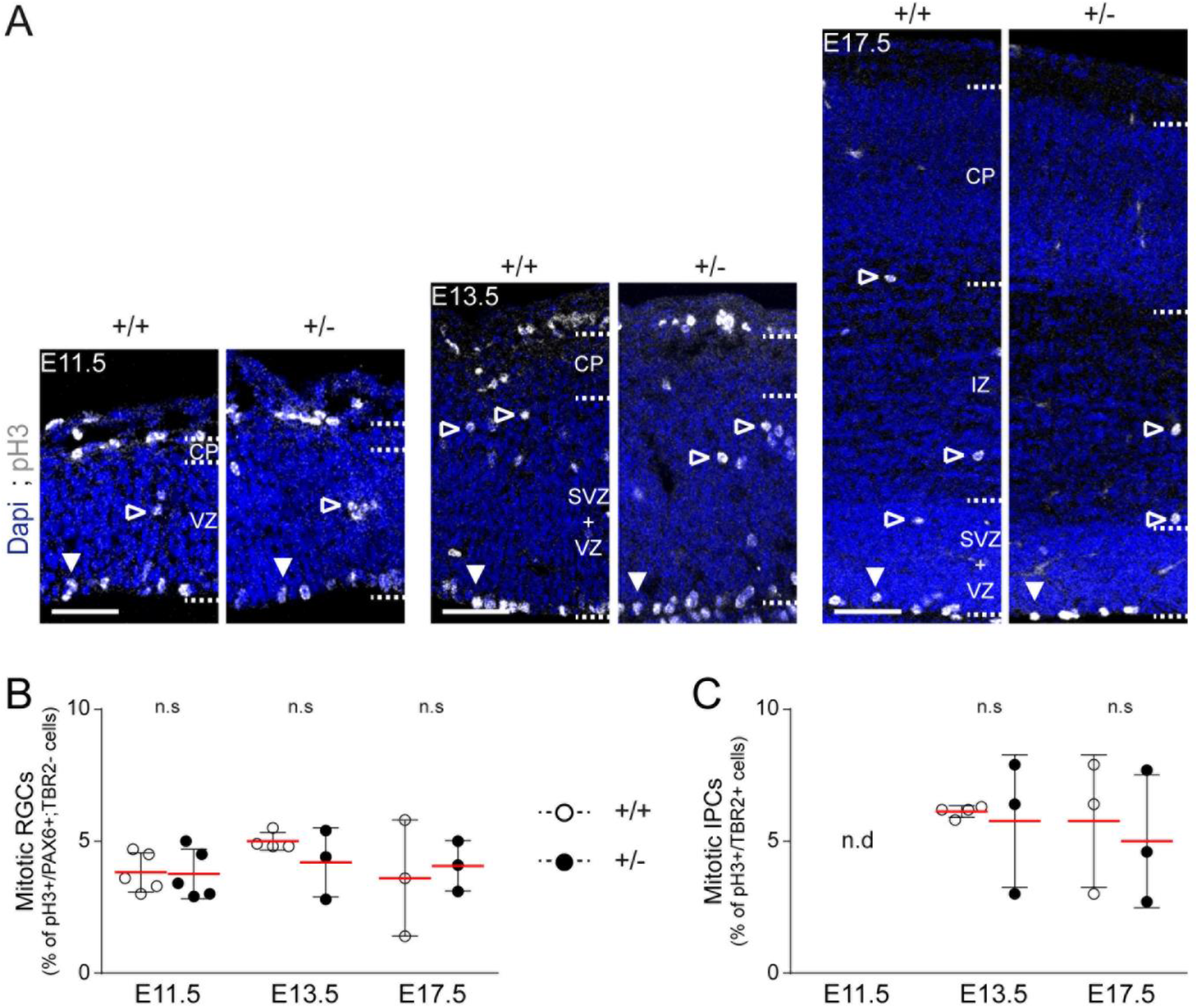
The mitotic index of RGCs and IPCs is not altered in the cerebral cortex of SmadNes mutant embryos. (A) Coronal sections showing apical (white arrowheads) and basal (black arrowheads) pH3+ mitoses in the developing cerebral cortex of SmadNes mutant embryos (*Smad1^wt/fl^;Smad5^wt/fl^;Nestin:Cre^+/0^*, +/−) and control littermates (*Smad1^wt/fl^;Smad5^wt/fl^;Nestin:Cre^0/0^*, +/+) at E11.5, E13.5 and E17.5. (B, C) Mean proportion ± s.d of (B) pH3+/PAX6+;TBR2-cells (mitotic RGCs) and (C) pH3+/TBR2+ cells (mitotic IPCs) quantified in a 100 μm-wide cortical area, obtained from 5, 4, 3 +/+ and 5, 3, 3 +/− embryos at E11.5, E13.5 and E17.5, respectively. Significance was assessed at each developmental stage with the two-sided unpaired t-test. ns: P>0.05; nd: not determined. Scale bars, 50 μm. CP: cortical plate, IZ: intermediate zone, SVZ: sub-ventricular zone, VZ: ventricular zone.

**Figure S7:**
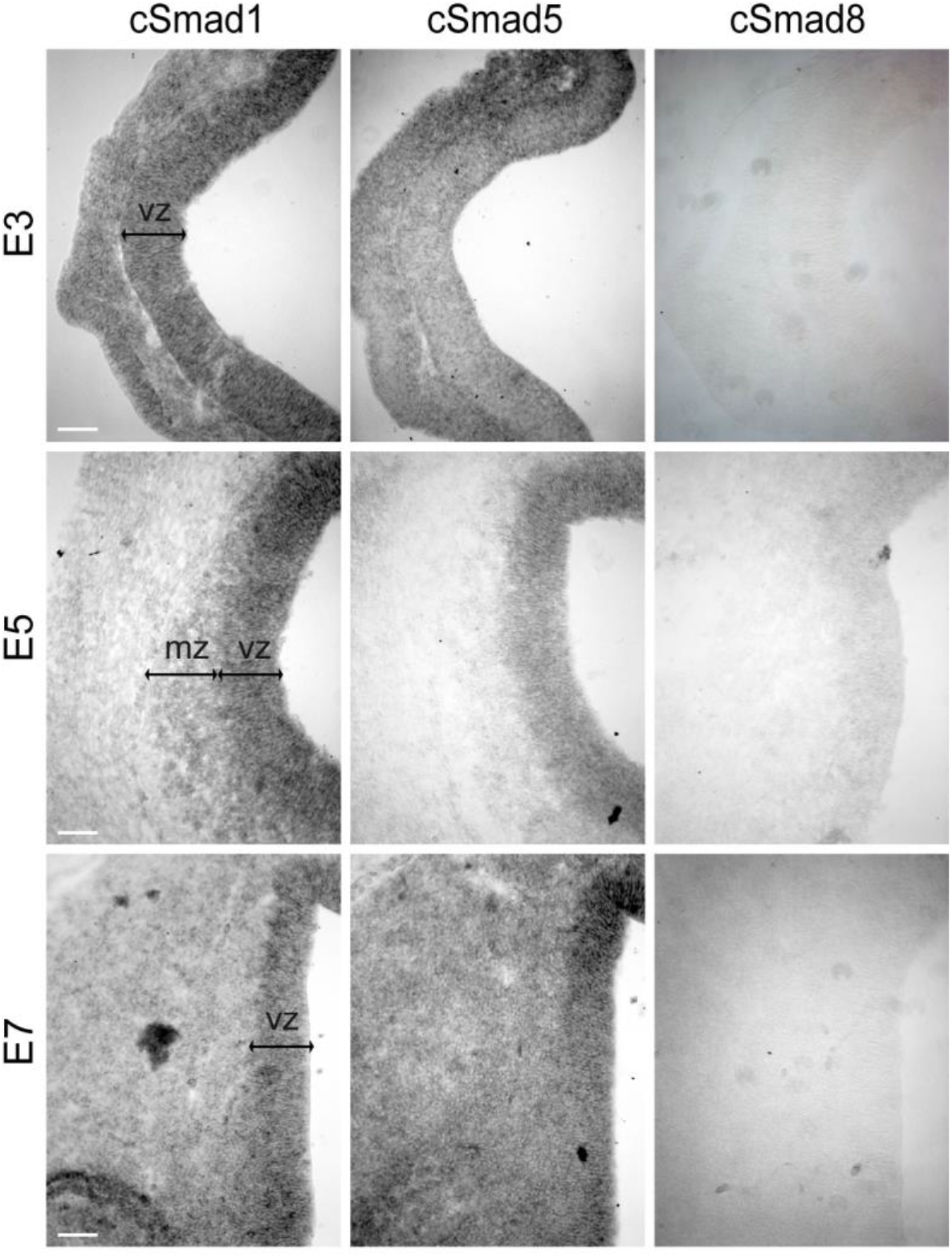
Expression of cSmad1/5/8 during chick cortical neurogenesis. Coronal sections showing *cSmad1, cSmad5 and cSmad8* transcripts detected by in situ hybridization in the developing cerebral cortex of E3, E5 and E7 chick embryos. Scale bars: 50 μm. VZ: ventricular zone, MZ: mantle zone.

**Figure S8:**
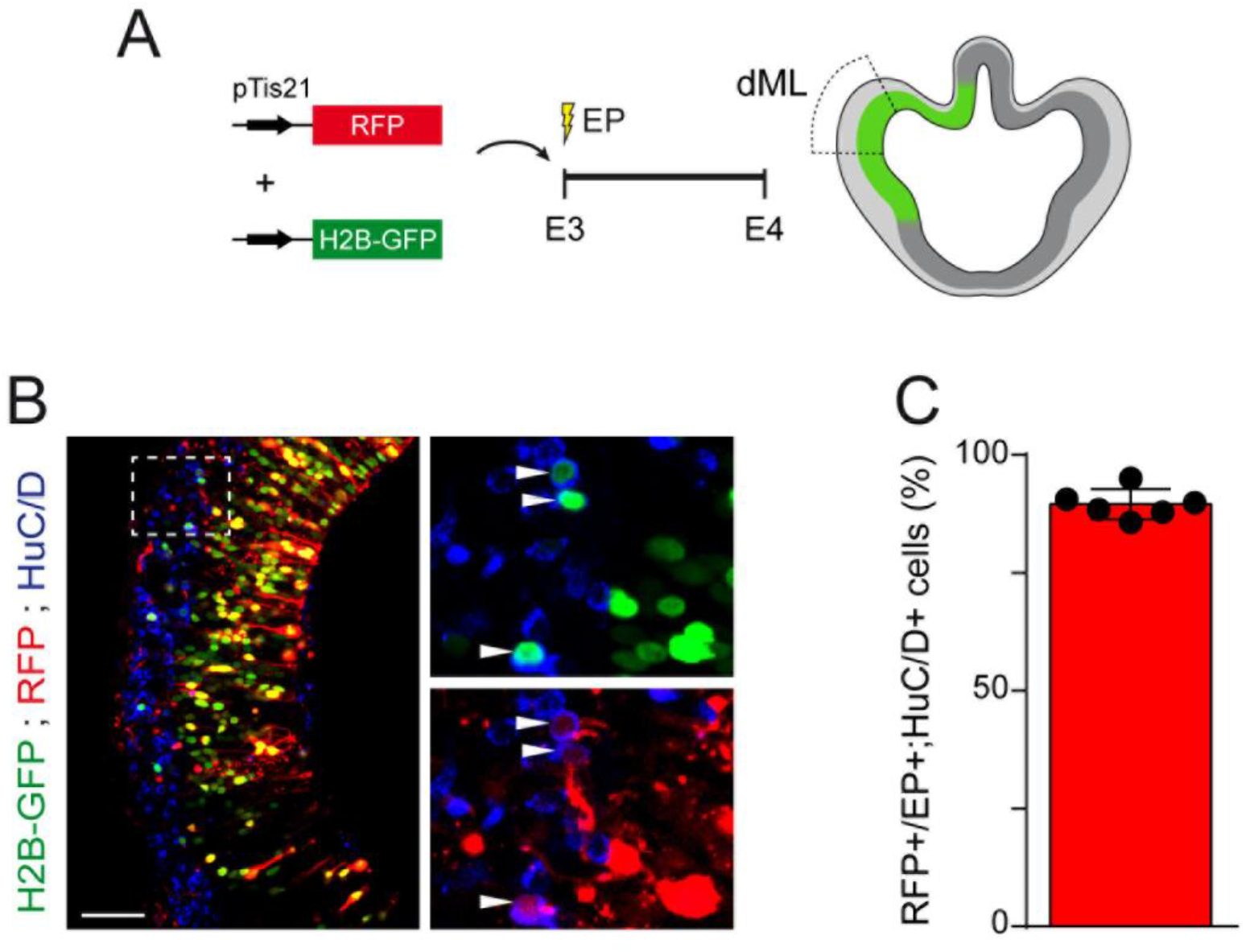
The activity of the pTis21:RFP reporter identifies neurogenic divisions during early chick cortical neurogenesis. (A) Representation of the methodology used to identify neurogenic divisions during early chick cortical neurogenesis. The pTis21:RFP reporter was co-electroporated together with a constitutively active H2B-GFP-producing plasmid, and its activity assessed in the dorsal medial-lateral (dML) cortical region 24 hours after in ovo electroporation (IOE) of the dorsal telencephalon of E3 chick embryos. (B) Representative image of pTis21:RFP+ cells among the electroporated (H2B-GFP+) cells that differentiated into HuC/D+ neurons 24 hours later and (C) its quantification, presenting the mean proportion of RFP+ cells among H2B-GFP+;HuC/D+ cells ± s.d, obtained from 6 electroporated embryos. Scale bars: 50 μm.

**Figure S9:**
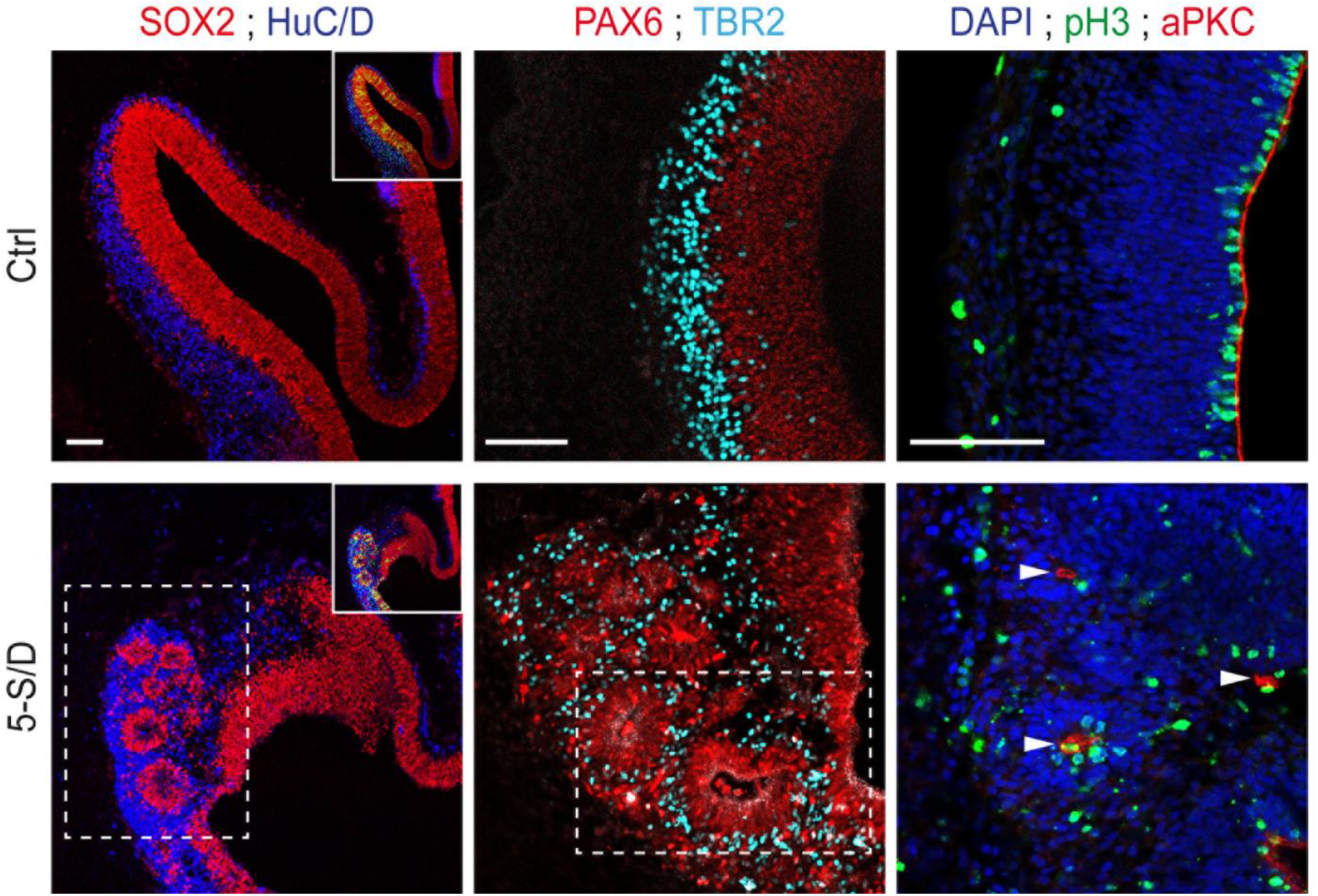
Increasing SMAD5 activity causes the abnormal generation of ectopic rosettes of cortical progenitors. Representative images showing that IOE of SMAD5-S/D can cause the abnormal generation of ectopic rosettes of SOX2+ cortical progenitors. The PAX6+ and TBR2+ cells forming these rosettes developed an ectopic apical-basal polarity, as revealed by immunostaining for the apical marker atypical PKC (aPKC, arrowheads), and divide mostly at the center of the rosettes (pH3+). This phenotype has been observed in the cerebral cortex of 25% (5 out of 20) of the embryos electroporated with SMAD5-SD. Scale bars: 50 μm.

**Figure S10:**
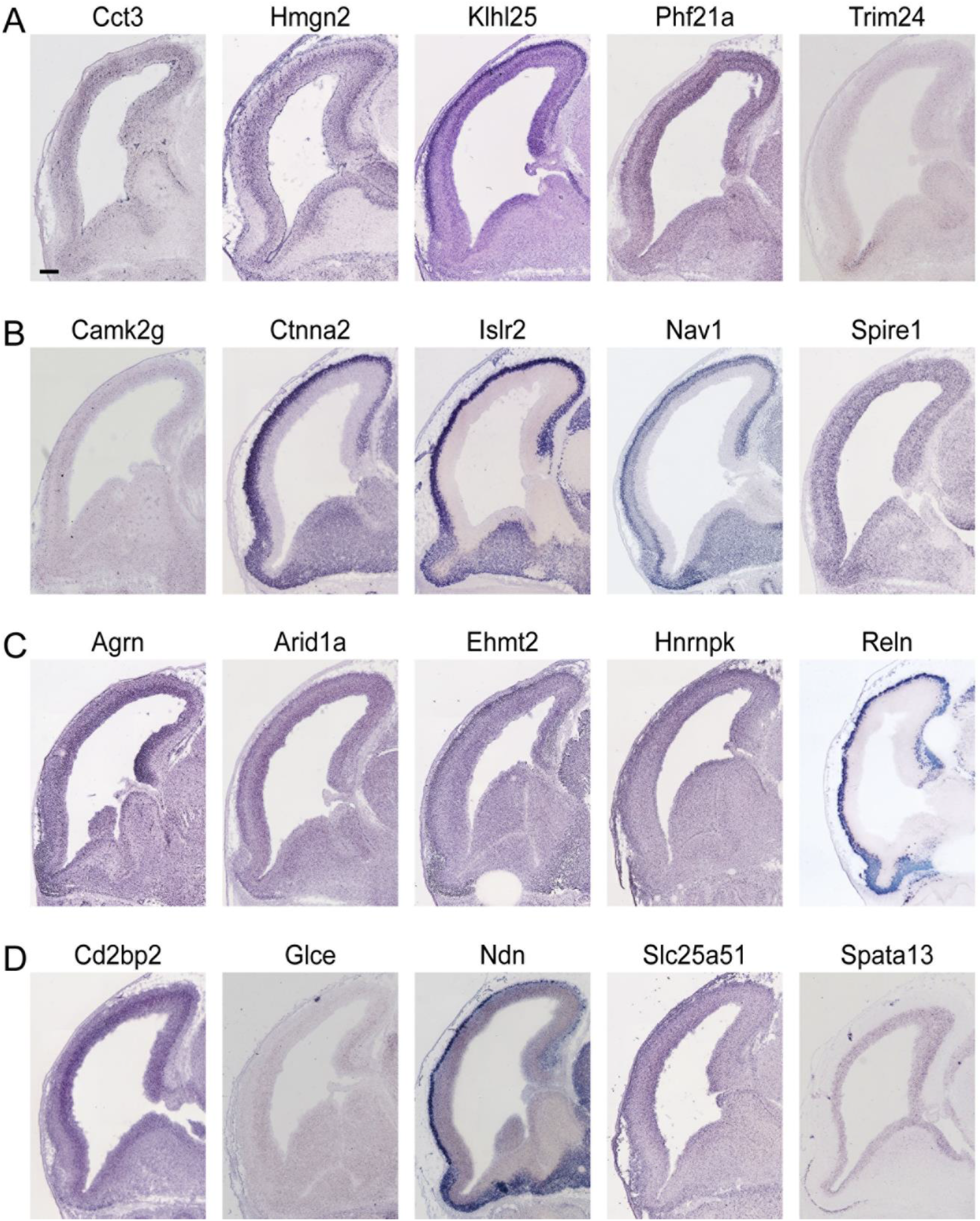
Gene expression pattern of RNAseq candidates in the mouse telencephalon. Sagittal sections of the developing mouse dorsal telencephalon at E14.5 showing the gene expression pattern of candidates identified in the RNA-seq, including (A-C) genes whose promoters are enriched in binding motifs for TEADs and retrieved in the GO term related to (A) cellular biosynthesis such as *Cct3,Hmgn2, Klhl25, Phf21a, Trim24,* (B) neurogenesis such as Camk2g, Ctnna2, Islr2, Nav1, Spire1, (C) both such as *Agrn, Arid1a, Ehmt2, Hnrnpk, Reln* and (D) genes not related to TEADs, such as *Cd2bp2, Glce, Ndn, Slc25a51, Spata13.* All the images were obtained from Genepaint (https://gp3.mpg.de). Scale bars: 50 μm.

**Figure S11:**
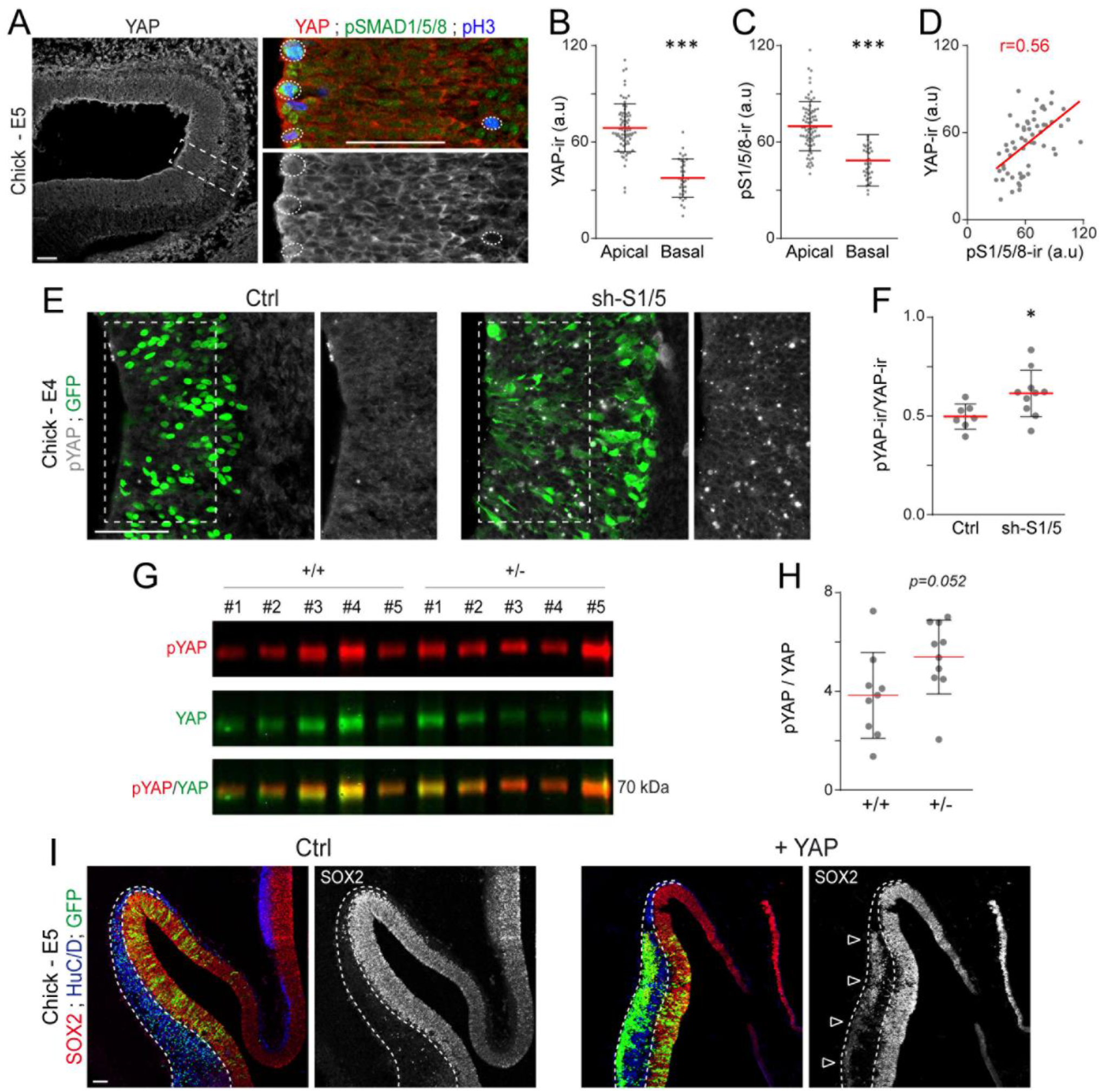
SMAD1/5 regulate early cortical neurogenesis through YAP. (A) Expression of YAP during chick corticogenesis, relative to SMAD1/5 activity (pSMAD1/5/8). (B,C) The mean intensity ± s.d of endogenous YAP (B) and pSMAD1/5/8 (C) immunoreactivity was quantified in 74 apical and 33 basal mitotic (pH3+) nuclei obtained from 3 chick E5 embryos. (D) The correlation between the intensity of endogenous pSMAD1/5/8 and YAP immunoreactivities was assessed by calculating the Pearson’s correlation coefficient r. (E) Immunostaining of the form of YAP phosphorylated at Serine 123 (pYAP) and (F) quantification of the mean ratio ± s.d of pYAP/YAP intensity obtained 24 hours after IOE with sh-S1/5 (10 sections from 3 embryos) or its control (7 sections from 3 embryos). (G) Western blot detection of the levels of pYAP and total YAP in extracts obtained from telencephalons ofE11.5 SmadNes mutant embryos (*Smad1^wt/fl^;Smad5^wt/fl^;Nestin:Cre^+/0^*, +/−) and their control littermates (*Smad1^wt/fl^;Smad5^wt/fl^;Nestin:Cre^0/0^*, +/+). (H) Quantification of the mean pYAP/YAP ratio ± s.d, obtained from 10 SmadNes mutant embryos (+/−) and 9 control littermates (+/+) derived from 2 different litters. (I) Immunostaining of the developing chick cerebral cortex for SOX2+ neural progenitors and HuC/D+ differentiating neurons 48 hours after IOE with a construct overexpressing the wild type form of YAP (YAP) or a control, showing that YAP overexpression can lead to a massive ectopic delamination of SOX2+ electroporated cells in the basal part of the mantle zone. Significance was assessed with the non-parametric Mann–Whitney test (B, C, F), the two-sided unpaired t-test (D, H).*P<0.05, ***P<0.001. Scale bars, 50 μm.

**Table S5:** List of primary antibodies used in this study.

| REAGENT or RESOURCE | SOURCE | IDENTIFIER |
| --- | --- | --- |
| Antibodies |  |  |
| anti-aPKC (mouse) | Santa Cruz Biotechnologies | Cat #SC-17781 (H-1); RRID:AB_628148 |
| anti-BrdU (rat) | AbD Serotec | Cat #MCA2060; RRID:AB_323427 |
| anti-APC clone CC1 (mouse) | Millipore | Cat #OP80; RRID:AB_2057371 |
| anti-CTIP2 (rat) | Abcam | Cat #ab18465; RRID:AB_2064130 |
| anti-CUX1 (rabbit) | Proteintech | Cat #11733-1-AP; RRID:AB_2086995 |
| anti-pHistone3 (rat) | Sigma Aldrich | Cat #H9908; RRID:AB_260096 |
| anti-HuC/D (mouse) | Life Technologies | Cat #A-21271; RRID:AB_221448 |
| anti-MEF2c (rabbit) | Abcam | Cat #ab64644; RRID:AB_2142861 |
| anti-NeuN (mouse) | Millipore | Cat #MAB377; RRID: AB_2298772 |
| anti-OLIG2 (rabbit) | Millipore | Cat #AB9610; RRID: AB_570666 |
| anti-PAX6 (mouse) | DSHB | Cat #pax6; RRID:AB_528427 |
| anti-PAX6 (rabbit) | Covance / Biolegend | Cat #901301; RRID:AB_2565003 |
| anti-Prominin1 (rat) | eBioscience | Cat #17-1331-81; RRID: AB_823120 |
| anti-SATB2 (mouse) | Abcam | Cat #ab51502; RRID:AB_882455 |
| anti-SMAD1 (rabbit) | Cell Signaling Technologies | Cat #6944S; RRID:AB_10858882 |
| anti-SMAD5 (rabbit) | Cell Signaling Technologies | Cat #12534S; RRID: n.d |
| anti-pSMAD1/5/8 (rabbit) | Cell Signaling Technologies | Cat #9511; RRID:AB_331671 |
| anti-SOX2 (rabbit) | Abcam | Cat #ab97959; RRID:AB_2341193 |
| anti-SOX9 (rabbit) | Kindly provided by Dr. M. Wegner | RRID:AB_2315360 |
| anti-SOX10 (goat) | R&D Systems | Cat #AF2864; RRID:AB_442208 |
| anti-TBR1 (rabbit) | Abcam | Cat #ab31940; RRID:AB_2200219 |
| anti-TBR2 (rabbit) | Abcam | Cat #ab23345; RRID:AB_778267 |
| anti-TUBULIN-b3 (Tuj1) (mouse) | Covance | Cat #MMS-435P; RRID:AB_2313773 |
| anti-Vinculin (mouse) | Sigma | Cat #V9131; RRID: AB_477629 |
| anti-YAP (mouse) | Santa Cruz Biotechnologies | Cat #sc-101199; RRID:AB_1131430 |
| anti-pYAP (rabbit) | Cell Signaling Technologies | Cat #4911; RRID:AB_2218913 |

